# Loss of tumour suppressor p53 rewires enhancer landscape and governs oncogenic progression

**DOI:** 10.1101/2025.07.28.667341

**Authors:** Harsha Rani, Dimple Notani, Vijayalakshmi Mahadevan

## Abstract

Mutations in tumour suppressor p53 confer enhanced metastasis and chemoresistance in colorectal cancer (CRC). Though the genetic events regulating CRC with p53 loss/mutation have been documented, the epigenetic events accompanying the loss of p53 have not been well understood. Epigenome based classification of CRC tumours has identified the active enhancer mark as a distinct marker for progression, however the role of the distal regulatory regions upon p53 loss in CRC remains to be established. This work investigates the influence of p53 loss on enhancer regulation in colorectal cancer cells. Genome wide profiling of active enhancer mark, H3K27ac in p53wt and p53-/- CRC cells reveal an overall gain of this mark around the promoters and intronic regions. These active enhancers show strong association with oncogenes and hallmark MYC and E2F targets suggesting an enhancer mediated regulation of MYC/E2F pathway governed by E2Fs, MAZ and PATZ1. Interestingly, we also observed a gain in oncogenic super enhancers mediated by E2Fs/KLFs accompanying loss of p53. The promoters of histone methyl transferases *EZH2 and SuV39H1* (E2F targets) show elevated levels of H3K27ac suggesting a novel epigenetic regulation of CRC around the promoters and distal regulatory regions. Our validation of these findings in p53 deficient colon cancer cohorts shows that the super enhancer associated genes align more to the CMS4 subtype and exhibit lower survivability. The observed cancer stemness and gain of oncogenic super enhancers with p53 loss presents a hitherto unexplored paradigm of enhancer mediated oncogenic progression which may be exploited for devising epigenetic therapy in p53-/- CRC patients.

**Significance:** Colorectal cancers (CRC) lose tumour-suppressor function and gain neomorphic functions with mutation/loss of p53. This work explores the epigenomic modulation of p53 null CRC cells by distal regulatory elements which has not been not clearly understood yet. We report a global increase in the active enhancer mark H3K27ac at active promoter and enhancer regions. We find that the gained enhancers/promoters are regulated by E2Fs/MAZ/PATZ1 which drive cancer stemness while the lost enhancers/promoters are regulated by tumour-suppressive IRFs. The activation of E2Fs correlates with elevated H3K27ac implying positive feedback driving E2F targets such as EZH2 and SuV39H1. The indirect activation of histone methyltransferases by p53 and the gain of oncogenic super-enhancers present a novel epigenetic regulatory paradigm which we also validated in p53 null CRC cohorts. These findings aid the design of epigenetic therapy for p53 deficient colorectal tumours.

## Introduction

Colorectal cancer (CRC) is the second leading cause of cancer deaths worldwide and the third most prevalent solid cancer (Bhandari et al., 2017). Colorectal cancer has been associated with three major pathways: Chromosomal Instability (CIN), Micro Satellite Instability (MSI) and CpG island Methylator Phenotype (CIMP) (Worthley et al., 2010).

International Cancer Consortia have further classified colorectal cancer into four consensus molecular subtypes (CMS) based on molecular signatures as CMS1 (microsatellite instability immune, 14%), hypermutated, microsatellite unstable and strong immune activation; CMS2 (canonical, 37%), epithelial, marked WNT and MYC signaling activation; CMS3 (metabolic, 13%), epithelial and evident metabolic dysregulation; and CMS4 (mesenchymal, 23%), prominent transforming growth factor–β activation, stromal invasion and angiogenesis (Guinney et al., 2015).

Recent work investigating prevalent somatic mutation patterns in CRC reported across 15 public datasets revealed mutations across 7 key genes with a mutational frequency of ≥ 10%. These genes include Tp53 (67 %), APC (66 %), KRAS (43 %), PIK3CA (18 %), FBXW7 (14 %), SMAD4 (14 %), and BRAF (10%). The combination of APC, Tp53 and KRAS has also been shown to confer the poorest survival in CRC (Su et al., 2024). Initial CMS classification has shown Tp53 mutations to be associated more with CMS2 and CMS4 subtypes (Guinney et al., 2015, Nzitakera et al., 2024) that exhibit aggressive cancer phenotypes. Missense mutations of known to deactivate the tumor suppressor function of Tp53 and activate gain of novel oncogenic functions including cell proliferation, antiapoptotic effects and metastatic formation and are therefore defined as the Gain of Function (GoF) mutations (Rivlin et al., 2011), (Muller & Vousden, 2013)

I*n vivo* and *in vitro* model systems have shown different phenotypes associated with distinct p53 mutations (Lowe et al., 1994,Hassin et al., 2022,Sabapathy & Lane, 2018, Baugh et al., 2017). The type of p53 mutation and the phenotype associated with each mutation thus become a crucial factor in determining the effectiveness of the therapy administered. Transcriptional reprogramming involving the activation of oncogenes and repression of tumour suppressor genes is a significant event relating to phenotype changes associated with cancer progression. Alterations in regulatory landscape mediated by transcription factors (TFs), cofactors, chromatin regulators, directly or indirectly bind to transcriptional regulatory elements, including promoters, enhancers and super-enhancers (SEs), thereby governing major reprogramming event in cancer progression.

Post translational modifications of histone are dysregulated across cancers and act as a bookmark in defining this regulatory landscape. (Karczmarski et al., 2014,Qin et al., 2020). These histone tags including H3K27ac, H3K4me3 are responsible for either the activation and repression of regulatory elements including promoters and enhancers which in turn control gene expression. Cancer cells acquire super-enhancer regions which provide a substrate for signaling pathways to regulate tumour suppressors and oncogenes like c-MYC and Bcl-2 and(Hnisz et al., 2015). Reprogramming of distal regulatory elements such as enhancers and super-enhancers are also associated with therapeutic resistance and cellular plasticity making them potential substrates for intervention in cancer therapy (Bi et al., 2020). Genome wide profiling of the active enhancer mark H3K27ac and the active epigenetic mark H3K4me3 in colorectal cancer tissues have identified a global increase of these marks at both the enhancers and promoters (Q. L. Li et al., 2021) compared to adjacent normal tissues suggesting an association of their elevated levels with cancer progression.

Enhancers are known to function as active (H3K27ac), primed (H3K4me1), closed or poised enhancers (H3K4me1 and H3K27me3) (Shlyueva et al., 2014). Comprehensive epigenomic comparison and characterization of colorectal adenocarcinomas, adenomas and normal tissues identified H3K27ac marked active enhancers as the most discriminatory epigenetic element, characterizing various histopathological stages of colorectal cancer. Epigenome-based classification (EpiC) has classified colon cancer into four different clusters: EpiC1, EpiC2, EpiC3 and EpiC4, with EpiC2 and EpiC4 enhancers to have significantly higher genome wide enrichment compared to the other two clusters. These EpiC clusters overlap with CMS classification highlighting the importance of p53 mutation in defining the high level of enhancers (Orouji et al., 2022)These observations suggest a combination therapy of MEKi plus BRDi as a general therapeutic strategy in colorectal cancer. Similar study has highlighted the epigenomic enhancer dysregulation that occurs in parallel to DNA mutations that occur at canonical proto-oncogenes and tumour suppressors during malignant transformation of colon crypt cells into CRC (Cohen et al., 2017).

Experiments over the last decade have suggested that GoF Tp53 mutants gain their potency by targeting cellular pathways regulating chromatin structure (Sammons et al., 2015), (Prives & Lowe, 2015)). Mutant p53 regulates a distinct set of histone methylases MLL1 and MLL2 and acetyl transferase MOZ, implying the involvement of novel epigenetic pathways that co-opt genetic pathways to promote cancer metastasis (Zhu et al., 2015). Since distinct p53 mutations associate with distinct phenotype, it becomes crucial to understand the regulatory mechanisms and the phenotypes associated with the mutation/loss of p53. Recent work (Chang et al., 2023) has established that p53 deletion enhances the tumour initiating potential of multiple myeloma cells besides promoting its self-renewal property and enhanced pan clinical drug resistance in patients harbouring deletion of 17p13.

The epigenetic landscape mediating cancer progression with p53 deletion/loss and their associated phenotypic changes have not been well established. Although studies have implicated elevated levels of the enhancer H3K27ac with cancer progression (Flebbe et al., 2019), epigenetic alterations accompanying loss in tumour suppressor p53 and the regulation of enhancer and super enhancer circuitry in modulating tumour progression in CRC has not been well established.

This work attempts to characterize regulatory elements through profiling of active enhancer mark H3K27ac in both p53 WT and p53 -/- CRC cells (HCT116 and HCT116 p53-/-) and understand the alterations in the epigenomic landscape and transcriptional circuitry driving oncogenic transformation upon p53 loss in CRC cells. We have performed ChIP sequencing to understand the genome wide enrichment of the enhancer mark H3K27ac and have identified regulators playing a direct role in activating oncogenes upon loss of p53 in CRC cells. On integrating the ChIP sequencing profile with the RNA-seq data, we observed that the loss of p53 is accompanied by a promoter-centric gain of H3K27ac correlating with increased expression of E2F and MYC targets. The sites that gained H3K27ac are mostly driven by E2F/SP1 like transcriptional circuitry while the lost sites accommodate interferon like transcription factors (IRFs).

Interestingly, we observed an indirect gain of H3K27ac around the promoters and enhancers of histone methyl transferases EZH2 and SuV39H1 which are known E2F targets with the gain of the enhancer mark correlating with enhanced expression of the methylases. We also report for the first time, a novel finding that the loss of p53 is accompanied by a gain in oncogenic super enhancers in colorectal cancer cells. The gained super enhancer regions are associated with various oncogenes mostly due to indirect effect of loss of p53. The observed network of epigenetic regulation and the altered regulatory transcriptional circuitry have also been validated in colorectal patient tumours with loss of p53. The tumours derived from the TCGA-COAD show increased expression of oncogenes associated with gained super-enhancers in p53 null tumours as compared to p53WT tumours. The increased oncogenic expression correlates with lower survivability in colon tumours. This observation adds a new and novel layer of epigenetic regulation of p53 loss in colorectal cancer and may be exploited for targeted epigenetic therapy in patients who show deletions in p53.

## Results

### Loss of p53 leads to a global increase in the active H3K27ac histone mark

In order to understand the genome wide enrichment of the enhancer mark upon the loss of p53, we performed a series of chromatin immunoprecipitation assays followed by high-throughput sequencing (ChIP-seq) for H3K27ac.

The ChIP sequencing yielded a total of 20,845 and 25,057 significant peaks (IDR (irreproducible discovery rate) <0.05) in p53 WT and p53-/- CRC cells respectively suggesting a gain of enhancer marks upon loss of p53. To further examine these differences at the genome scale, we performed differential binding analysis on this active histone mark using normalized H3K27ac signal values around the merged peak sets and identified 18149 sites to be gained, 1851 sites to be lost while 6375 sites unaltered upon p53 loss (Log2FC |0.5| and p-val < 0.05) (Fig 1A-B, FigS1A-B), again suggesting that pre-existing H3K27ac-enhancers further gained the mark whereas, several regions that did not exhibit H3K27ac in WT also showed gain upon p53 loss. The high levels of H3K27ac levels observed in these experiments indicate robust transcriptional state of these cells upon p53 deletion.

**Fig 1.**
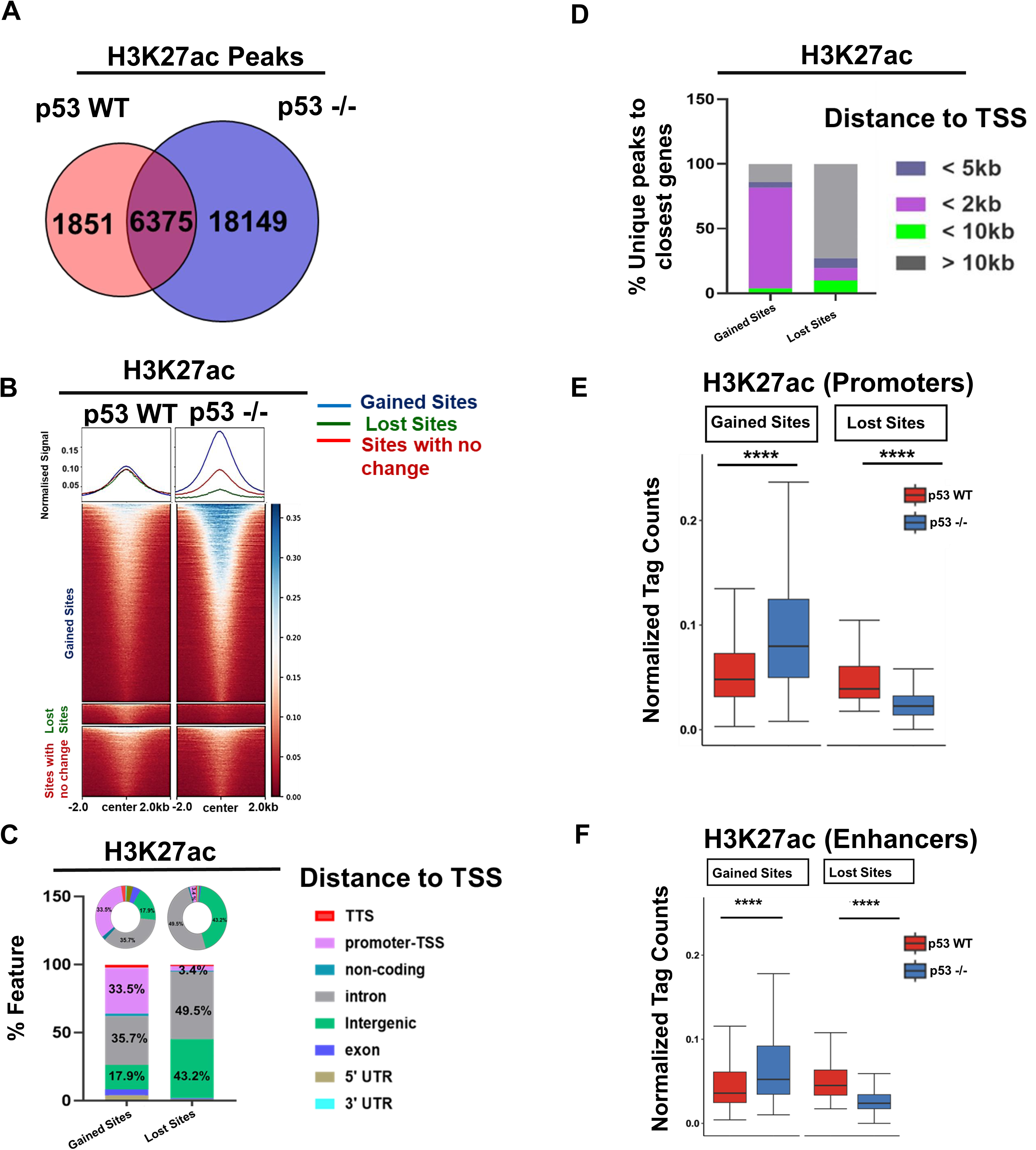
Loss of p53 leads to a global increase in enhancer mark H3K27ac in colorectal cancer cells. A. Venn Diagram representing the overlap of H3K27ac ChIP-seq peaks between p53WT and p53-/- CRC cells. The loss of p53 is accompanied by gain in H3K27ac binding sites. B. The signal intensity profile of H3K27ac centered at +/- 2kb of gained sites (blue), lost sites (green) and sites with no change (red) upon p53 loss in CRC cells. The profile represents an overall gain in the enhancer mark H3K27ac upon loss of p53 C. Genome wide distribution of gained and lost H3K27ac sites upon p53 loss in CRC cells represented as stacked bar plot and pie chart. The profile represents increased occupancy of H3K27ac in the promoter region (pink) followed by the intronic regions (grey) upon loss of p53 D. Percentage of unique genes closest to the peak of gained and lost sites upon p53 loss in CRC cells represented as a stacked bar plot. The profile reflects that the gained sites show the highest enrichment at the promoters within 2kb (<2kb) distance from the TSS while the lost H3K27ac sites show enrichment at distal regions(>10kb). E. Box plot representing normalized tag counts of H3K27ac at gained and lost promoters (+/-2Kb of TSS) upon p53 loss in CRC cells [p53 WT (red) and p53-/- (blue)] (Wilcoxon-Rank test was performed for the statistical analysis, p value> 0.05, ns; p value<=0.05, *; p value<=0.01, **; p value<=0.001, ***). The promoter sites which gained H3K27ac signals are accompanied by significant gain in levels of H3K27ac while the promoter sites which lost H3K27ac signals are accompanied by significant loss in the levels of H3K27ac F. Box plot representing normalized tag counts of H3K27ac at gained and lost enhancers (away from +/-2Kb of TSS) upon p53 loss in CRC cells [p53 WT (red) and p53-/- (blue)] (Wilcoxon-Rank test was performed for the statistical analysis, p value> 0.05, ns; p value<=0.05, *; p value<=0.01, **; p value<=0.001, ***). The enhancer sites which gained H3K27ac signals are accompanied by significant gain in levels of H3K27ac while the enhancer sites which lost H3K27ac signals are accompanied by significant loss in the levels of H3K27ac.

We next analyzed the genome wide distribution of gained and lost H3K27ac peaks using HOMER (Heinz et al., 2010) and observed a clear over representation of promoter-proximal (+/-2Kb) and intronic regions among the gained H3K27ac sites while the sites losing H3K27ac were found to be clustered more around the intronic and intergenic regions (Fig1E-F). On comparison, we observed that only 3.4% of the lost H3K27ac sites, but 33.4% of gained H3K27ac sites overlapped with promoters (Fig1E-F). In contrast, 43.2% of lost H3K27ac sites were located in distal intergenic regions as compared to 17.9% of gained H3K27ac sites. Interestingly, this suggests that WT p53-dependent gene activation accompanied by H3K27ac are likely to be regulated more by distal p53-binding events. Though the observed promoter and intronic gain in H3K27ac correlating with the increase in active mark around both the promoters and enhancers in various cancers (Q. L. Li et al., 2021, Huang et al., 2021) has been reported, the increase in enhancer marks due to the loss of p53 in colorectal cancer cells is an interesting and novel layer of observation from this work. To further characterize these regulatory elements, we defined the active promoter peaks as those that appear within 2 kb of transcription start sites and the remaining peaks (away from 2kb of TSS) as active enhancer peaks. Out of the total 18149 gained H3K27ac sites, 6771 (37.30%) were associated with distal regulatory enhancer sites (away from ± 2kb from TSS region) defined as gained enhancers and 11378 sites (62.69%) were identified as promoters, defined as gained promoters (FigS1E-F). Similarly, out of the total 1851 lost H3K27ac sites, 1736 sites were associated with distal regulatory regions, defined as lost enhancers (93.8%) and 115 sites (6.1%) were identified as promoters, defined as lost promoters. The H3K27ac signal levels strongly establish these findings (Fig1G and H).

### Loss of p53 upregulates oncogenic programs in CRC by activation of MYC and G2M checkpoints targets

To gain insight into how the transcriptional reprogramming observed upon loss of p53 in colorectal cancer cells leads to oncogenic phenotypes, we carried out RNA-sequencing to understand the global gene expression profiles of p53 WT and p53-/- CRC cells. We observed 5443 significantly differentially expressed genes (with p-value less than 0.05.), out of which 2,529 were upregulated (log FC>=0.5) and 2,521 genes were downregulated (log FC<=-0.5) upon p53 loss as compared to p53 WT CRC cells (Fig2A, S2A). Out of these differentially expressed genes, a majority of the genes regulating the p53 signalling pathway were found to be downregulated (103 genes were found to be downregulated while 64 were upregulated) (Fig2A, S2B). We then performed preRanked gene set enrichment analysis (GSEA) based on the gene expression (higher to lower) values. This GSEA of differentially expressed genes (pvalue <= 0.05) showed upregulation of oncogenic pathways such as the MYC transcriptional signature and the G2M checkpoint gene sets (Fig. 2B, S2C), suggesting that WT p53 represses MYC and G2M checkpoint target genes. The differential gene expression profile also showed a negative enrichment score for p53 signalling pathway (Fig 2B). The enrichment of RNA polymerase activity and mRNA splicing observed through gene ontology analysis also (Fig 2C) suggest an increased transcriptional and post transcriptional activity in CRC upon p53 loss correlating with increased levels of H3K27ac.

**Fig 2.**
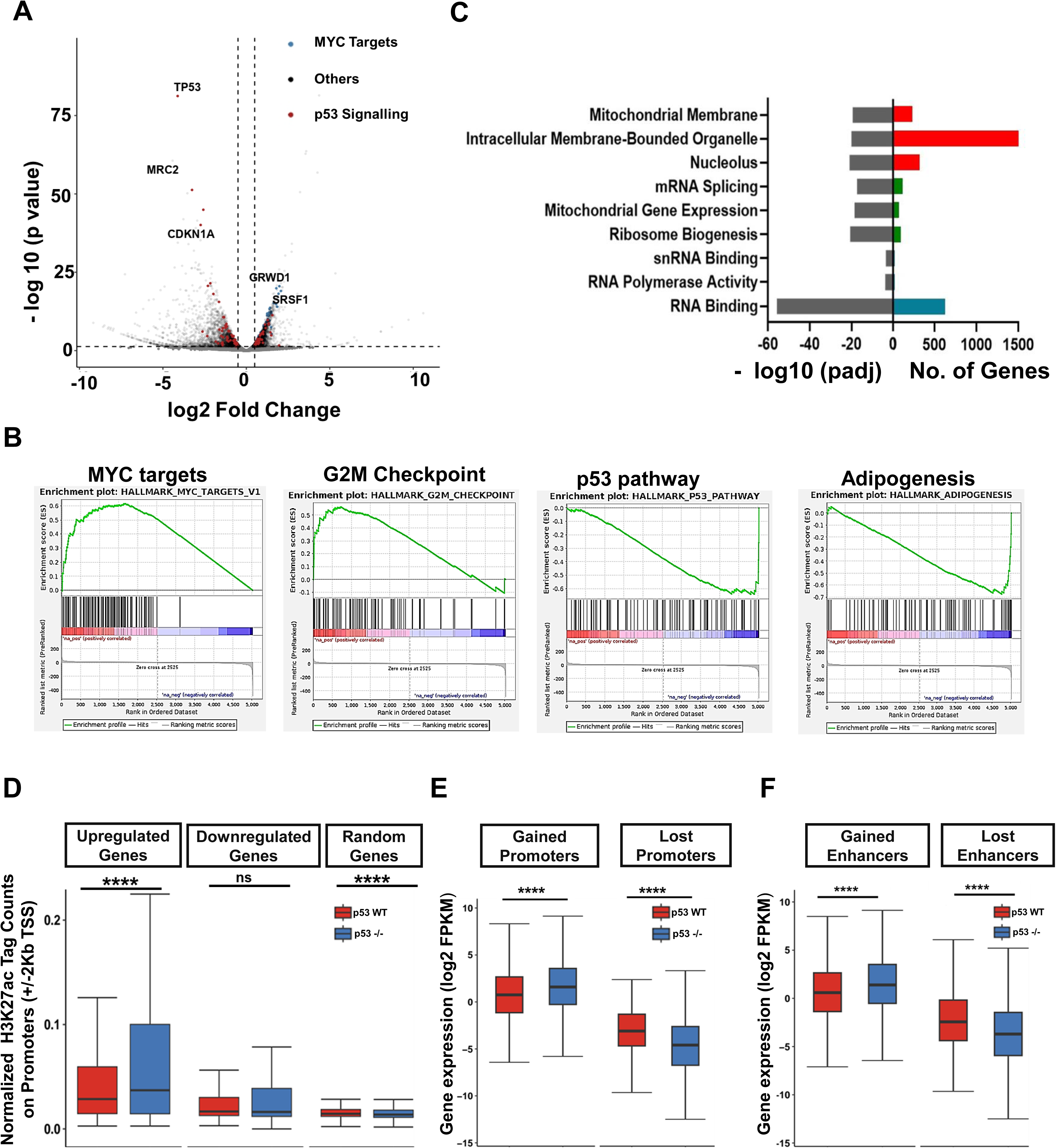
Loss of p53 potentiates epigenetic reprogramming at transcriptionally altered genes. **A.** Volcano plot representing differential gene expression profile in p53 WT Vs p53 -/- CRC cells [p value <=0.05 and log2FC|0.5|]. Highlighted dots represent significantly expressed genes involved in p53 signalling (red) and MYC targets (blue). Loss of p53 shows dysregulation of genes involved in p53 signalling and MYC targets. **B.** Positive enrichment score of MYC targets and G2M-checkpoint upon p53 loss as revealed through preRanked Gene set enrichment analysis (GSEA) on differential enrichment of gene signatures with p value <=0.05. Negative enrichment of adipogenesis and p53 pathway upon p53 loss in CRC cells with p value <=0.05. Loss of p53 activates MYC and G2M and downregulates p53 signalling pathway. **C.** Gene Ontology analysis representing enriched cellular component, molecular function and biological process upon loss of p53 (p value <=0.05). It can be observed that the genes upregulated with the loss of p53 are involved in RNA polymerase activity and mRNA splicing. **D.** Box plot representing distribution of H3K27ac counts at the promoters of upregulated, downregulated and random genes (t-test was performed for the statistical analysis, p value > 0.05, ns; p value<=0.05, *; p value<=0.01, **; p value<=0.001, ***). The promoters of upregulated genes are accompanied by a significant increase in the levels of H3K27ac. **E.** Box plot representing distribution of gene expression (Log2FPKM) at the gained and lost promoters (t-test was performed for the statistical analysis, p value> 0.05, ns; p value<=0.05, *; p value<=0.01, **; p value<=0.001, ***) in p53 WT (red) and p53-/- (blue) CRC cells. Genes associated with gained promoters correspond to a significant increase in expression profile while the genes associated with lost promoters correspond to a significant decrease in expression profile upon loss of p53 **F.** Box plot representing the distribution of gene expression (Log2FPKM) at the putative gained and lost enhancers (t-test was performed for the statistical analysis, p value> 0.05, ns; p value<=0.05, *; p value<=0.01, **; p value<=0.001, ***) in p53 WT (red) and p53-/- (blue) CRC cells. Genes associated with gained enhancers correspond with significant increase in expression profile while the genes associated with lost enhancers correspond with significant decrease in expression profile upon loss of p53.

In order to understand the functional implications of enhancer gain on the expression profile of the upregulated, downregulated and a set of random genes, we identified the promoters of differentially expressed genes with higher change in expression profile (with an absolute log FC cutoff as 1 and p-value as less than 0.05) and random genes for which no change in gene expression levels). We observed that the promoters of upregulated genes correspond significantly (p value <=0.05) to the elevation in H3K27ac in p53-/- CRC cells as compared to p53-WT while in contrast the promoters of downregulated genes also showed an insignificant and marginal increase in H3K27ac levels. The random genes showed no alteration in the H3K27ac levels upon p53 loss (Fig 2D). This suggests that the increased H3K27ac correlates significantly with the highly expressed upregulated genes.

To explore functional consequences of the alterations in H3K27ac levels at the identified regulatory elements upon p53 loss, we associated gained and lost sites around promoters with the nearest genes using bedtools closest (Quinlan & Hall, 2010). When the genes associated with the promoters were further overlapped with the set of differentially expressed genes, we found that the genes associated with the gained promoters correspond to the significant increase in expression in p53-/- CRC cells (Fig 2E) as compared to p53-WT CRC cells, while the genes associated with the lost promoters correspond to the significant decrease in expression profile in p53-/- CRC as compared to p53 WT CRC cells (Fig 2E). This is consistent with the number of genes identified to be significantly upregulated [log2FC >= 0.5 and pvalue <=0.05] corresponding to the increase in H3K27ac levels at the gained promoters (2312 genes) (Fig S2D), while 230 genes [log2FC <=-0.5 and pvalue <=0.05] to be downregulated corresponding to the decrease in H3K27ac signals at the lost promoters (Fig S2D). Gene set enrichment analysis of the genes associated with gained H3K27ac at the promoters revealed a positive enrichment score for MYC and E2F targets (FigS2F) suggesting that the epigenetic reprogramming of the enhancer mark upon p53 loss drives colon cancer towards metastasis and stemness by modulating E2F and MYC pathways.

To further characterize the impact or functional correlation of gained and lost H3K27ac at enhancers, we associated each enhancer to genes within 1Mb distance using the GREAT algorithm (McLean et al., 2010) and overlapped them with the differentially expressed genes. We found that the gained enhancers correspond significantly to the increase in gene expression profile in p53-/- CRC cells as compared to p53-WT CRCs while the lost enhancers correspond significantly to the decrease in gene expression profiles (Fig 2H). This is consistent with number of downregulated genes associated with the lost enhancers (387 genes, pvalue <=0.05 and logFC>0.5) (Fig S2E) and the number of upregulated genes associated with gained enhancers (807 genes, pvalue <=0.05 and logFC <=-0.5). Furthermore, gene set enrichment analysis revealed a negative enrichment score of Interferon gamma response around the lost enhancer sites (Fig S2G). Loss of IFNγ receptor or its response has been shown to drive colorectal cancer progression and has been reported in patients with colorectal cancer (Wang et al., 2015, Du et al., 2022). We hence hypothesize that loss of p53 promotes cancer progression through suppression of interferon gamma response and enhancing MYC and E2F targets through alteration in H3K27ac levels at enhancers and promoters respectively.

### Transcriptional circuitries governed by E2Fs and IRFs at the regulatory regions upon p53 loss in CRC cells

To identify the principal transcriptional factors associated with the gained and lost H3K27ac sites upon p53 loss, we performed differential motif analysis using X-Streme (MEME-Suite) around the gained sites using lost sites as a control and *vice - versa*. We observed that upon the loss of p53, GC rich motifs corresponding to E2Fs, MAZ and PATZ1 were enriched around the gained H3K27ac sites (Fig3A, S3A), while those around the lost sites showed higher enrichment of AT rich motifs corresponding to GATA, MEFs, PIT1, Tp53 and IRFs transcription factors (Fig 3B, S3B).

**Fig 3.**
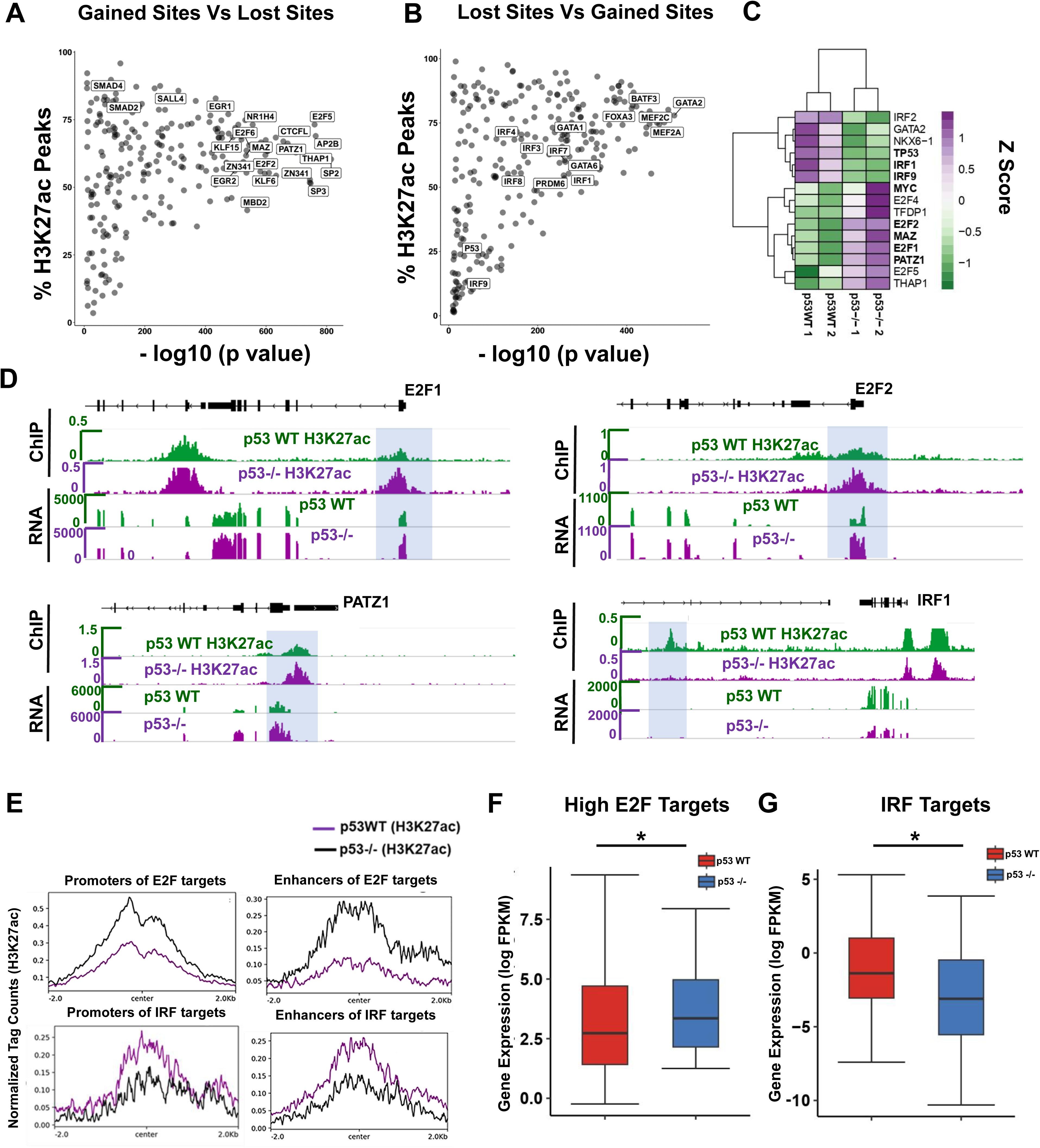
Transcriptional circuitry governed by E2Fs and IRFs at the regulatory regions upon p53 loss in CRC cells. **A.** Scatter plot representing differential motif analysis at the gained H3K27ac as compared to lost H3K27ac sites, with Y axis representing the percentage of H3K27ac sites. The highlighted regions represent the transcription factors associated with higher significance value upon p53 loss. Transcription factors like E2Fs and KLFs were associated with higher percentage of gained H3K27ac sites. **B.** Scatter plot representing differential motif analysis at the lost H3K27ac as compared to gained H3K27ac sites, with y axis representing percentage of H3K27ac sites. The highlighted regions represent the key transcription factors associated with higher significance value upon p53 loss. Transcription factors like IRFs and MEFs were associated with higher percentage of lost H3K27ac sites. **C.** Heat map representing differential expression profile of the identified transcription factor in p53 WT (green) and p53-/- (purple) CRC cells (p value<=0.05, logFC|0.5|). Upregulation of the transcription factors including E2Fs, MAZ and PATZ1 while downregulation of IRF1, Tp53 and GATA2 upon loss of p53. **D.** IGV screenshot representing H3K27ac profiles in p53 WT (green) and p53-/- (purple) CRC cells integrated with the RNA tracks at the promoters of E2F1, E2F2, PATZ1 and IRF1. Increased expression of E2F1 and E2F2 are associated with gain of H3K27ac around the promoters. E2F2, E2F1 and PATZ1 are upregulated upon p53 loss. Decreased expression of IRF1 was found to be associated with loss of H3K27ac around the putative enhancers. **E.** Signal intensity profile at the promoters and enhancers of E2F targets (upper panel) while at the promoters and enhancers of IRF targets (lower panel). Promoters and enhancers of the identified E2F targets associated with increased H3K27ac levels, while the promoters and enhancers of the identified IRF targets associate with decreased H3K27ac levels. **F.** Box plot representing distribution of expression profiles of gene targets of High E2F expressing genes and IRF (student t-test was performed for the statistical analysis, p value> 0.05, ns; p value<=0.05, *; p value<=0.01, **; p value<=0.001, ***) in p53 WT (red) and p53-/- (blue) CRC cells. Expression profiles of genes involved in E2F correlate with a significant increase in expression profile, while IRF targets correlate with significant decrease in expression profile.

In order to understand whether distinct transcriptional regulators identified in gained and lost sites correlates with changes in gene expression profiles, we overlapped the TFs identified at the distinct sites with the gene expression profiles. Interestingly, we found that most of E2Fs (E2F1, E2F2 and E2F4) were upregulated upon p53 loss along with MAZ and PATZ1 while IRFs were found to be downregulated upon p53 loss suggesting a distinct transcriptional network regulating the events due to p53 loss (Fig 3C).

The expression profiles of E2Fs, MAZ and PATZ1 correlated with increased H3K27ac at the TSS region while decreased expression of IRF1 correlated with decreased H3K27ac sites at their putative enhancers (Fig 3D, S3D). E2F mediated target expression and cellular proliferation have been reported to be regulated by a p53 dependent negative feedback loop (Timmers et al., 2007) suggesting that loss of p53 activates the expression profile of E2F mediated targets and the increased transcriptional activity of E2Fs is mostly driven by an autoregulatory feedback mechanism associated with gained H3K27ac upon p53 loss.

Therefore, we next tried to understand the nature of the transcriptional circuitry governed by E2Fs at the gained promoters and enhancers upon p53 loss in CRC cells. We divided E2Fs target genes into two groups, high and low expressing E2F targets, and found that high E2F expressing groups showed a significant gain in expression profile as compared to low expressing E2F targets (FigS3D). We identified 95 upregulated E2F target genes to be associated with gained H3K27ac sites around the promoters and 33 upregulated genes to be associated with gained H2K27ac sites around the enhancer regions (Fig 3E, 3F). Interestingly, among these upregulated E2F targets genes, which correlate with increase in H3K27ac levels at both the promoters and enhancer regions are the histone methyl transferases EZH2 and SuV39H1 (Fig S3E).

We also found that the lost H3K27ac sites were associated with various interferon regulatory factors. To further understand the transcriptional circuitry regulating the Interferon receptor factors and interferon signalling pathway in CRC cells upon p53 loss, we examined the expression profiles of their target genes associated with lost H3K27ac sites at promoters and enhancers respectively (Fig3E-F). We found that most of them are significantly downregulated around the lost enhancer sites upon p53 loss. These include interleukin-15(IL15) which plays an important role in supporting the maturation of NK cells and apoptotic genes including CASP4, suggesting that WT p53 interacts with various IRFs in maintaining the tumour suppressive environment in CRC.

### Loss of p53 causes gain in super enhancers around oncogenes

Super enhancers are defined as those regulatory elements within the genome which span several kilobase pair length and is enriched with various epigenetic modifications including H3K27ac, coactivators and transcription factors(Whyte et al., 2013). The master transcription factors (Oct4, Sox2 and Nanog) form unusual enhancer domains at most genes that control the pluripotent state. Super-enhancers differ from typical enhancers in size, transcription factor density and content, their ability to activate transcription and in their sensitivity to perturbation types (Whyte et al., 2013).

Synergistic therapy involving BET inhibitor treatment JQ1 and cisplatin was shown to enhance the survival of ovarian cancer bearing mice in orthotopic models by suppressing ALDH activity and abrogating BRD4-mediated ALDH1A1 expression through a super-enhancer element (Yokoyama et al., 2016). CRC-related SEs is known to promote the transcriptional expression of targeted oncogenes including MYC targets (Whyte et al., 2013, Hnisz et al., 2015, Ying et al., 2020). Hence to explore if the loss of p53 mediates gain in super enhancers in colorectal cancer cells, we first identified super enhancer formation in p53WT and p53-/- CRC cells. Using HOMER software, the enhancer peaks within 12.5kb of each other were stitched together and the super enhancer associated peak regions were then ranked based on their signal intensity (Whyte et al., 2013).

We found a total of 729 super enhancers associated with p53 WT while 893 were associated with p53-/- CRC cells (Fig4A). In order to understand the differential enrichment of the identified super enhancers upon p53 loss, we computed the normalized H3K27ac signals around the merged super enhancer set using bedmap and identified 747 super enhancers to be gained (Log FC>0.5) and 19 super enhancers to be lost with a logFC <=-0.5 upon p53 loss (Fig4B). This suggests that the loss of p53 directly or indirectly induces super enhancer formation in CRC cells. Further, we quantified the number of H3K27ac peaks/gained super enhancer associated with p53-/- CRC cells and p53 WT CRC cells. We found that gained super enhancers harboured greater number of enhancer peaks from p53-/- CRC cells as compared to p53WT CRC cells (Fig 4C). This suggests that super enhancers gaining H3K27ac upon p53 loss have a greater number of active typical enhancer peaks associated within 12.5kb region as compared to those in cells expressing wild type p53. This implies that the distribution of the number of enhancer peaks varies in both p53WT and p53-/- CRC cells, with more enhancer peaks associated with p53-/- CRC cells. In order to correlate the alteration in H3K27ac levels around gained and lost super enhancers, we first looked into the average normalized tag counts of H3K27ac around the gained and lost super enhancer regions scaled to 5kb region. We observed that the identified gained and lost super enhancers regions correlate with alterations in H3K27ac levels (Fig 4D), that is gained super enhancers were associated with higher H3K27ac signal in p53-/- CRC as compared to p53WT CRC cells while the lost super enhancers were associated with lower H3K27ac signals in p53-/- CRC cells as compared to p53WT CRC cells.

**Fig 4.**
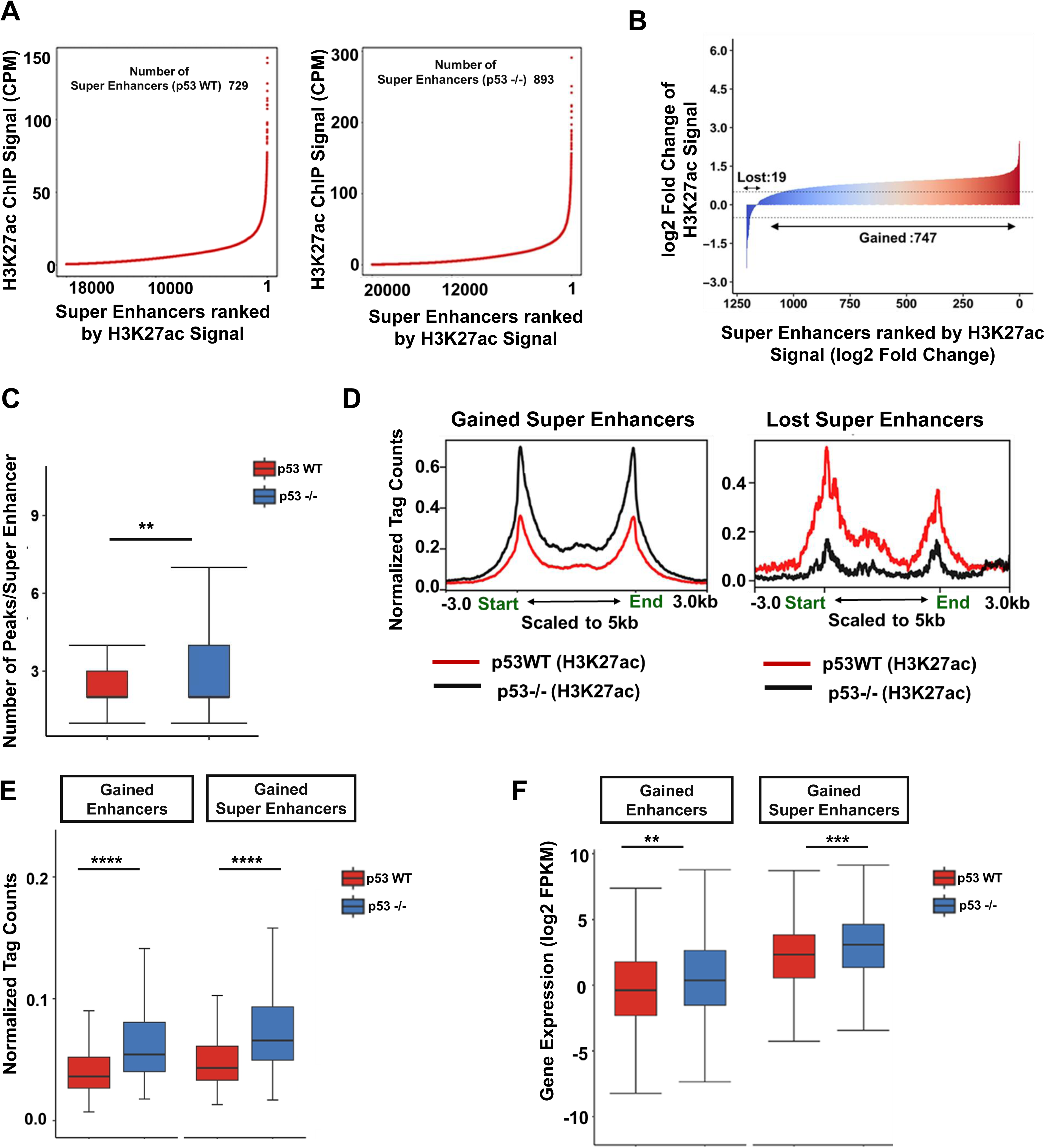
Loss of p53 induces gain in super-enhancers. **A.** Ranked plots of super enhancers (SEs) in the p53WT (right) and p53-/- CRC cells (left) obtained by increasing H3K27ac signal (normalized by the ratio to the highest H3K27ac reads of the enhancer). The values on the axes represent the ranks of enhancers by H3K27ac signal. Loss of p53 is associated with higher number of super- enhancers. **B.** Differential analysis for super-enhancers. All genomic regions containing a super- enhancer in p53WT and p53-/- CRC cells were ranked by log2 fold change in H3K27ac signal (p53WT Vs p53-/-). The x-axis shows log2 fold change in H3K27ac signals. Super- enhancers with log2 fold change| 0.5| were colored red or blue. Loss of p53 associates with gained super-enhancer regions. **C.** Box plot representing the number of H3K27ac peaks in p53 WT and p53 -/- per gained super enhancer regions (t-test was performed for the statistical analysis, p value > 0.05, ns; p value<=0.05, *; p value<=0.01, **; p value<=0.001, ***) in p53 WT and p53 -/- CRC cells. Super-enhancers with loss of p53 is accompanied by a greater number of typical enhancers as compared to p53 WT. **D.** H3K27ac profiles of p53 WT (red) and p53 -/- (black) CRC cells at the identified gained and lost super enhancers. Plots represent enrichment ± 3 kb around the gained and lost sites scaled to 5kb region. Gained super enhancers correlate with increased H3K27ac levels while lost super enhancers correlate with decreased H3K27ac levels. **E.** Box plot representing distribution of H3K27ac signal intensity profile around gained typical enhancers and gained super enhancers upon loss of p53(Wilcoxon-rank test was performed for the statistical analysis, p value > 0.05, ns; p value<=0.05,*; p value<=0.01,**; p value<=0.001,***) in p53 WT(red) and p53-/- (blue) CRC cells . Super- enhancers are associated with significantly higher H3K27ac levels as compared to typical enhancers. **F.** Box plot representing gene expression distribution profile (log2 FPKM) at typical gained enhancers and super enhancers (t-test was performed for the statistical analysis, p value> 0.05, ns; p value<=0.05, *; p value<=0.01, **; p value<=0.001, ***) in p53 WT (red) and p53-/-(blue) CRC cells. Expression of genes associated with gained super- enhancers are significantly higher as compared to typical enhancers.

We quantified and compared the levels of H3K27ac around the typical gained enhancers and super enhancers and found that the gained super enhancers were associated with higher H3K27ac signals as compared to typical gained enhancers (Fig 4E), while lost super enhancers were associated with greater loss in H3K27ac signals as compared to typical lost enhancer regions (FigS4A). In order to understand the functional implications of the identified gained and lost super enhancers, each of the super enhancer regions were associated with genes within 1Mb distance using GREAT algorithm. Integration with the differential gene expression profile further revealed that gained super enhancers were associated with higher gene expression in p53-/- CRC cells as compared to typical enhancers (Fig 4E) while the gene expression associated with lost super enhancers were associated with lower expression as compared to typical lost enhancers (Fig S4B). To further understand the functional role of the differentially expressed genes associated with gained super enhancers, we performed GSEA and observed a similar positive enrichment of hallmark gene sets including MYC and E2F (Fig S4C). This suggests that the loss of p53 induces more super enhancer formation around genes involved in oncogenic signals mainly by influencing MYC and E2F targets.

ChIP sequencing-based studies have established the overlapping binding sites of MYC and E2F and their common target genes in embryonic stem cell population (Chen et al., 2008). Furthermore, it has been shown that E2Fs and MYC act synergistically to control the S- G2 transcriptional program required for normal cell divisions and in maintaining crypt-villus integrity in Rb deficient mice model system (Liu et al., 2015). However, the regulation by E2Fs/MyC with the loss of p53 has not been established. Our observation therefore implies a novel regulatory network induced by loss of p53 that drives cancer progression through gain in super enhancers around E2Fs and MyC targets genes (Fig S4C, S4). In order to under to further understand the distinct transcriptional circuitry driving super enhancer mediated oncogenesis, we performed differential motif analysis around gained super enhancer regions using lost super enhancers as a control. We observed the enrichment of Krüppel-type zinc-finger transcription factor ZNF281, ZBT14, SP1, TAF1 like motifs around the gained super enhancer regions (Fig S4E). It has been established that activation of p53 indirectly represses ZNF281 through miR-34-a. Furthermore, ectopic expression of ZNF281 induces metastasis in a SNAIL dependent manner (Hahn et al., 2013). In this work, we found the enrichment of bromodomain-containing protein TAF1, a subunit of general transcription factor TFIID, which initiates preinitiation complex formation and cellular transcription. Biochemical approaches have further suggested interaction of MYC amino acids residues interact with TBP in the presence of the amino-terminal domain 1 of TBP-associated factor 1 (TAF1^TAND1^), This suggests that the loss of p53 drives MYC and E2F targets by enrichment of Kruppel like factors and TAF1 around the gained super enhancers.

We also observed a similar enrichment of tumour suppressors, Interferon regulatory factor (IRF1) and Tp53(Fig S4F) around the lost super enhancer regions. The presence of tumour suppressors including p53, IRFs around the lost super enhancers suggests that WTp53 has a distal regulatory effect in maintaining the tumour suppressive environment. This is affected through functioning with various other tumour suppressive transcription factors including various interferon regulatory factors. We therefore hypothesise that the gained H3K27ac sites and activation of oncogenes upon p53 loss is an indirect effect of p53 while lost H3K27ac sites and downregulation of tumour suppressive genes are mostly due to direct effect of p53.

Oncogenic super enhancers are identified as those super enhancers which are associated with genes involved in oncogenic progression. To further explore the different oncogenes associated super enhancers, we annotated the genes associated with gained super enhancers upon p53 loss. Out of the total 218 upregulated genes (log2FC>=0.5) associated with gained super enhancers, we found that 117 genes (53.6%) are involved in oncogenesis and metastatic progression in colon cancer. The top 20 such genes are represented in Fig 5A. An integration of the ChIP seq data of H3K27ac with the RNA sequencing data of these upregulated genes revealed that the super enhancers associated with KRT17 show higher levels of H3K27ac (Fig 5B). Upregulation of KRT17 is associated with poor prognosis in colon cancer patients and its overexpression has been shown to promote cancer invasion and metastasis through regulation of the Wnt/βcatenin pathway (Ji et al., 2021). *Invivo* chromatin immunoprecipitation analysis of p53 on a rat model system has identified two p53-binding sites in the promoter region of KRT17 and that the binding of p53 operates as a direct Krt17 repressor (Liao et al., 2016). This correlates with our observation that the loss of p53 upregulates KRT17 and this upregulation is mediated by gain in H3K27ac levels at both the promoters and super enhancer regions as shown in Fig 5B. Similarly, we found gain in H3K27ac levels around super enhancers of GADD45B, PRMT5 and EIF4G1. PRMT5 has been shown to regulate cell cycle progression and CRC cell growth mainly via activation of Akt (Yan et al., 2021) and its interaction with MCM7 (X. Li et al., 2021) and inhibition of PRMT5 using GSK3326595 activates p53 signalling pathway via the induction of alternative splicing of MDM4 and has been shown to play an essential role for cell survival through the regulation of eIF4E expression and p53 translation.

**Fig 5.**
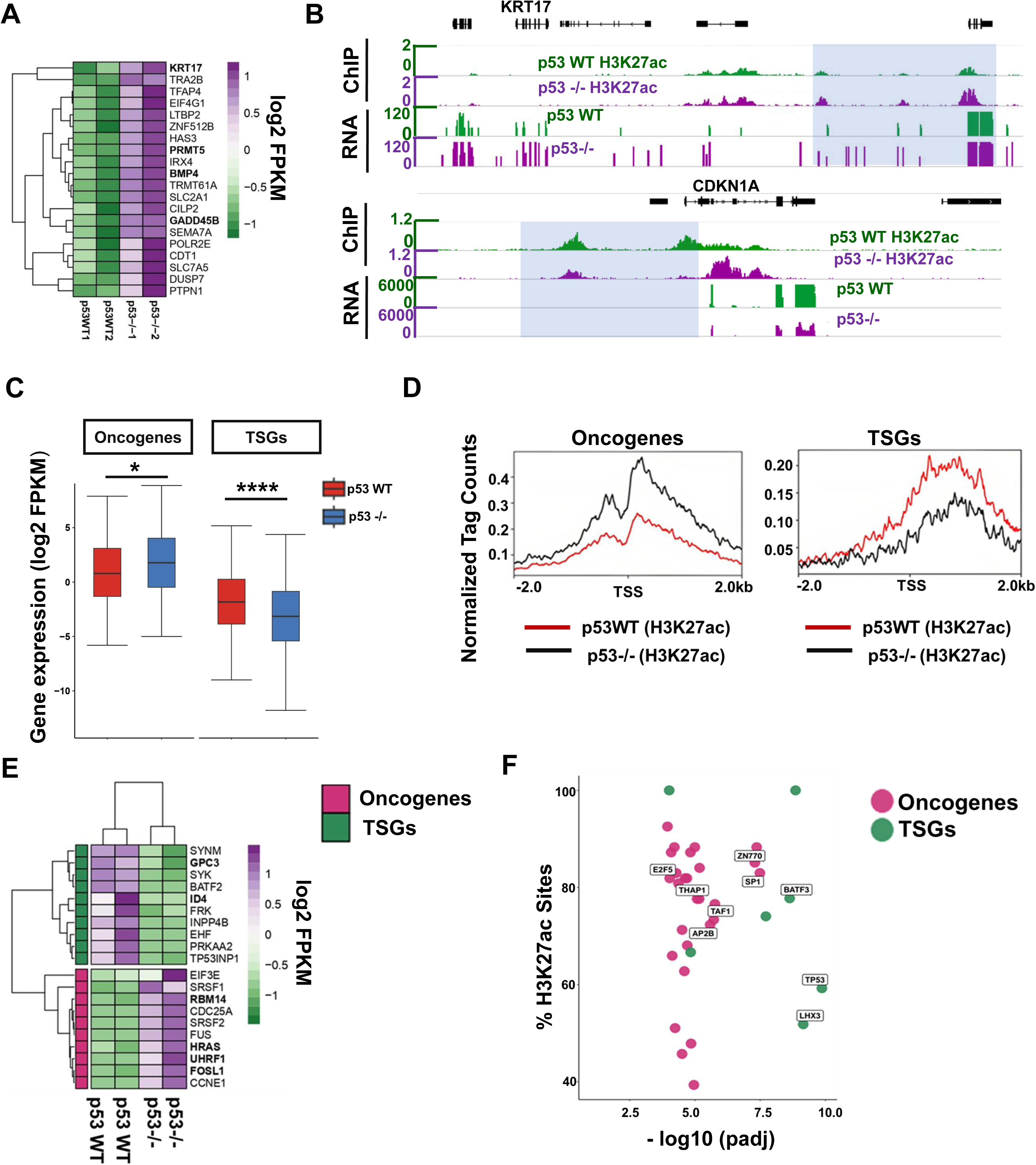
Dysregulation in the expression profile of tumor suppressors and oncogenes upon p53 loss in CRC cells. **A.** Heatmap representing top 20 upregulated oncogenes associated with gained super- enhancers in p53-/- CRC cells (p value<=0.05 and log2FC>=0.5). Gained super- enhancer associated oncogenes including KRT17, PRMT5, BMP4 and GADD45B are upregulated upon loss of p53. **B.** IGV screenshot representing H3K27ac profiles in p53 WT (green) and p53-/- (purple) CRC cells integrated with the RNA tracks around the super enhancers of KRT17 and CDKN1A. Increased expression of KRT17 correlates with gained of H3K27ac around the super-enhancers, while the decreased expression of CNKN2A correlates with loss of H3K27ac at the super-enhancers. **C.** Box plot representing gene expression distribution profiles of oncogenes and TSGs in p53 WT (red) and p53-/- (blue) CRC cells. Loss of p53 is accompanied with significant gain in the expression profile of oncogenes and significant decrease in the expression profile of tumour suppressors. **D.** H3K27ac profiles of p53 WT (red) and p53-/- (black) CRCs at the promoter region of differentially regulated oncogenes (upper panel) and TSGs (lower panel). Plots represent enrichment ± 0.5 kb around the peak center. Promoters of elevated expression of oncogenes correlates with elevated levels of H3K27ac while those of tumour suppressors correlates with decrease in the levels of H3K27ac. **E.** Heat map resenting top 10 downregulated TSGs(green) and top10 upregulated oncogenes(pink) upon p53 loss in CRC cells. Tumour suppressors including ID4 and GPC3 are downregulated upon loss of p53 while oncogenes including RBM14 and FOSL1 are upregulated upon loss of p53. **F.** Scatter plot representing differential enrichment of TF around the promoters of differentially expressed oncogenes(pink) and tumor suppressor genes(green) showing correlating with alterations in H3K27ac levels. E2F like transcription factorsare enriched at the promoters of oncogenes while Tp53 and IRFs enriched at the promoters of tumour suppressor genes.

### Direct regulation of tumor suppressors and indirect regulation of oncogenes by WTp53

The observation that the gained super enhancers accompany elevated oncogene expression upon p53 loss further led us to explore the alteration in expression profiles of various oncogenes and tumor suppressor genes across the CRC genome. We screened for the expression profiles of various known oncogenes and tumor suppressor genes (TSGs) known to be differentially regulated in colon cancer. Interestingly, we found that the loss of p53 correlates with the increase in expression profiles of various oncogenes and reduced tumor suppressor expression (Fig5C). To understand the epigenetic role in regulating the expression of various oncogenes and TSGs upon p53 loss we, associated each of these genes with the altered profiles of the enhancer mark H3K27ac. We identified around 116 oncogenes associated with gained H3K27ac around their promoters and 27 tumor suppressor genes associated with lost H3K27ac sites around their promoters (Fig5D), out of which top10 upregulated oncogenes and top10 downregulated genes associated with alterations in H3K27ac levels are represented as a heat map in Fig5E. This suggests that the loss of p53 directly or indirectly alters the regulatory landscape around various oncogenes and tumor suppressor genes to drive cancer progression.

To understand the transcriptional circuitry regulating the expression of tumor suppressors and oncogenes, we performed a differential motif analysis around the promoters of differentially regulated oncogenes and tumor suppressor genes correlating with alterations in H3K27ac levels. We found enrichment of p53 like transcription factor around the promoters of TSGs and E2Fs and SP1/KLFs around the promoters of oncogenes (Fig. 5F), suggesting that WT p53 exerts a direct influence in controlling the expression of tumor suppressors while indirectly controlling the expression of various oncogenes by activation of E2Fs/SP1. It has been proposed that WT p53 functions as an indirect repressor and a direct activator and that WT p53 regulates the baseline expression of various tumor suppressors ((Pappas et al., 2017)). This further correlates with our observation that oncogenes activated upon p53 loss are mostly driven by E2Fs/SP1/KLF like transcriptional circuitry as revealed through differential motif analysis while the repressed tumor suppressor genes are mostly driven by p53 and other tumor suppressors including IRFs which help maintain epithelial identity in CRC cells. This establishes that the loss of p53 drives the indirect activation of oncogenes and a direct repression of various TSGs. The indirect repression of E2Fs/SP1/KLF could be mainly through p21 mediated or noncoding miRNAs as previously described.

It is known that various stress signals and DNA damage activate the tumor suppressor p53, by disrupting the MDM2 and p53 complex, further triggering transcriptional activation of various downstream target genes (Hafner et al., 2019, Koo et al., 2022). In order to identify the binding sites of p53 around the overall gained and lost H3K27ac sites, we performed ChIP sequencing of p53 (naïve) on p53WT CRC cells and used the open source nutlin activated p53 ChIP sequencing done on p53WT HCT116 CRC cells. We identified 7339 peaks after macs2 peak calling (Fig6A, FigS6A). To understand the genomic distribution of p53 in these cells, we annotated p53 peak sets using HOMER. We observed that p53 peaks are mostly at intergenic regions (41.50%) and intronic regions (42.05%) and only 9.22% were enriched at the promoters suggesting that p53 has a distal regulatory effect in maintaining tumour suppressive environment (Fig6B). We analysed the p53 associated regions (+/-5kb) around the total gained and lost sites besides those with no changes. Out of the total 18,149 gained H3K27ac sites, we identified 1385 H3K27ac sites (7.6%) to be associated with p53. While, out of 1851 lost H3K27ac sites, 557(30.04%) sites were found to be associated with p53 and out of 6375 no change sites/persistent sites, 671 (10.52%) sites associated p53 (Fig6C). Furthermore, p53 signal strongly correlates with H3K27ac sites around the lost sites (Fig6B, FigS6B). A similar trend is observed with naïve p53 signals around gained, lost and no change sites (FigS6A). These observations suggest that WTp53 associates strongly with H3K27ac sites in p53 WT CRCs as compared to p53-/- CRC cells again suggesting a more distal regulatory effect.

**Fig 6.**
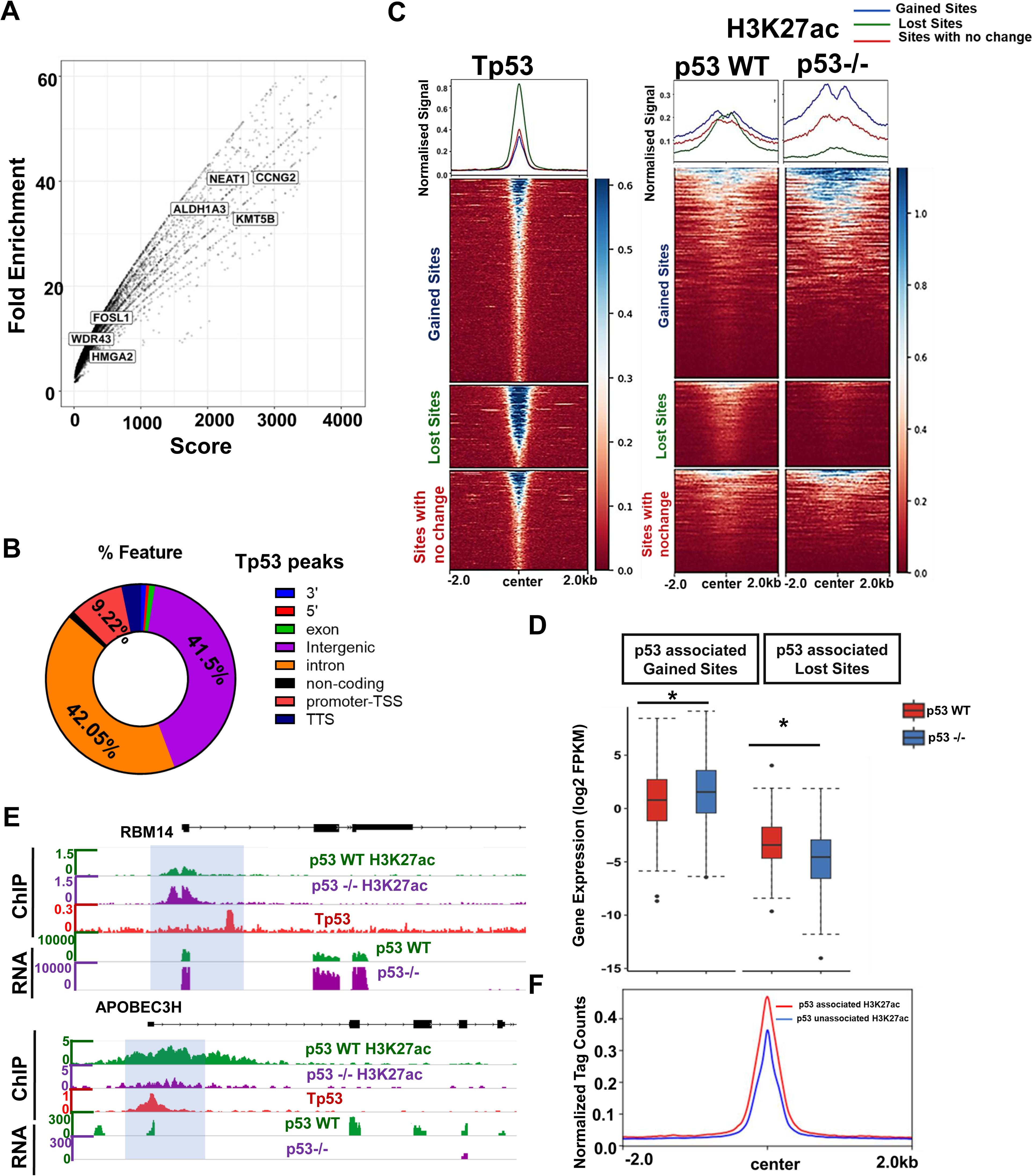
Direct regulation of tumour suppressors by wild type p53. **A.** Scatter plot of Tp53 ChIP seq (MACS2 output) with x-axis representing peak score and Y-axis representing fold enrichment of the p53 peaks. Higher enrichment score of activated p53 peaks are found near stemness related markers including ALDH1A3 **B.** Pie-chart showing the genomic wide distribution of the observed Tp53 peaks. Genome wide distribution of Tp53 peaks around the intronic region(orange) and distal intergenic region(purple) **C.** Heat map representing normalized p53 signal around p53 bound gained, lost and no change sites (left panel) and H3K27ac signals around the p53 bound gained, lost and no change sites (right panel) +/-2kb from the center of the peaks. **D.** Box plot representing distribution of gene expression (Log2FPKM) around p53 bound gained and lost sites (t-test was performed for the statistical analysis, pvalue> 0.05, ns; p value<=0.05, *; p value<=0.01, **; p value<=0.001, ***) in p53 WT(red) and p53-/- (blue) CRC cells. p53 associated gained H3K27ac sites correlates with significant gain in expression profile, while p53 associated lost H3K27ac sites correlates with significant loss in expression profile. **E.** IGV snapshots showing the alterations in H3K27ac, Tp53 binding sites and gene expression around APOBEC3H and RBM14. Decreased expression of APOBEC3H correlates with loss of H3K27ac and Tp53 binding sites. APOBEC3H was downregulated upon p53 loss. Increased expression of RBM14 correlates with gain of H3K27ac and Tp53 binding sites. RBM14 was upregulated upon p53 loss. **F.** p53 signal profile at the identified sites associated with H3K27ac region (Bound)(red) and not associated with H3K27ac(blue). Plots represent enrichment ± 2 kb around the binding site center. Tp53 signal is stronger with H3K27ac associated sites as compared to unassociated H3K27ac sites.

To understand the functional impact of p53 associated regions around gained and lost H3K27ac sites through gene expression profile, we associated each of these identified regions with genes (using bedtools closest) and analysed their overall expression profile. As expected, p53 regions associated with gained H3K27ac correlating with higher expression profiles, while the p53 regions associated with lost H3K27ac sites correlated with lower gene expression levels (Fig6D). We identified 135 upregulated genes with p53 associated gained H3K27ac sites and 49 downregulated genes with p53 associated lost H3K27ac sites. The p53 associated gained H3K27ac sites correlated with higher expression profiles of key oncogenes and stemness genes including RBM14 (Fig6E) and ALDH1A3 (FigS6C) upon p53 loss, while the lost H3K27ac sites correlated with lower expression of tumour suppressors and epithelial genes including APOBEC3H and ANK1(Fig6E and S6C).

Since only 2613 of the identified p53 peaks out of 7339 p53 binding sites were associated with genome wide active enhancer H3K27ac marks, we compared the p53 signals that were not associated with these H3K27ac sites (4726 peaks) in either of the two CRC cells. We compared the p53 signal around the p53 peaks associated with H3K27ac sites (p53 associated) with those that are not associated with H3K27ac sites (p53 non associated/unbound sites) and observed that p53 signals were higher in sites associated with H3K27ac sites as compared to the unbound sites (Fig6F, FigS6D). This clearly suggests that WT p53 associates strongly with the active enhancer mark H3K27ac and plays a direct activating role in maintaining the tumor suppressive environment in CRC cells.

### Oncogenic Super enhancer genes correlate with higher expression profiles in p53null CRC patients and lower survivability

In order to validate and correlate the gene signatures associated with altered regulatory landscape upon p53 loss in CRC cells with patient tumours we used the sequencing data of colon cancer patients from the TCGA-COAD database. We first stratified colon tumours based on the different somatic mutation profiles of Tp53. We divided them into two cohorts, p53 null tumors(N=61) defined as those which has truncating mutations in Tp53 including frame shift insertion, deletion, splice site variation and non-sense mutation which leads to deletion in full length Tp53 and p53WT tumors which do not bear any mutation in p53(N=61) (Fig7A). We first, correlated the expression profile of p53 in these patients and found that expression level of Tp53 was significantly downregulated in p53 null CRC patients as compared to p53 WT CRC patients (Fig 7B).

**Fig 7.**
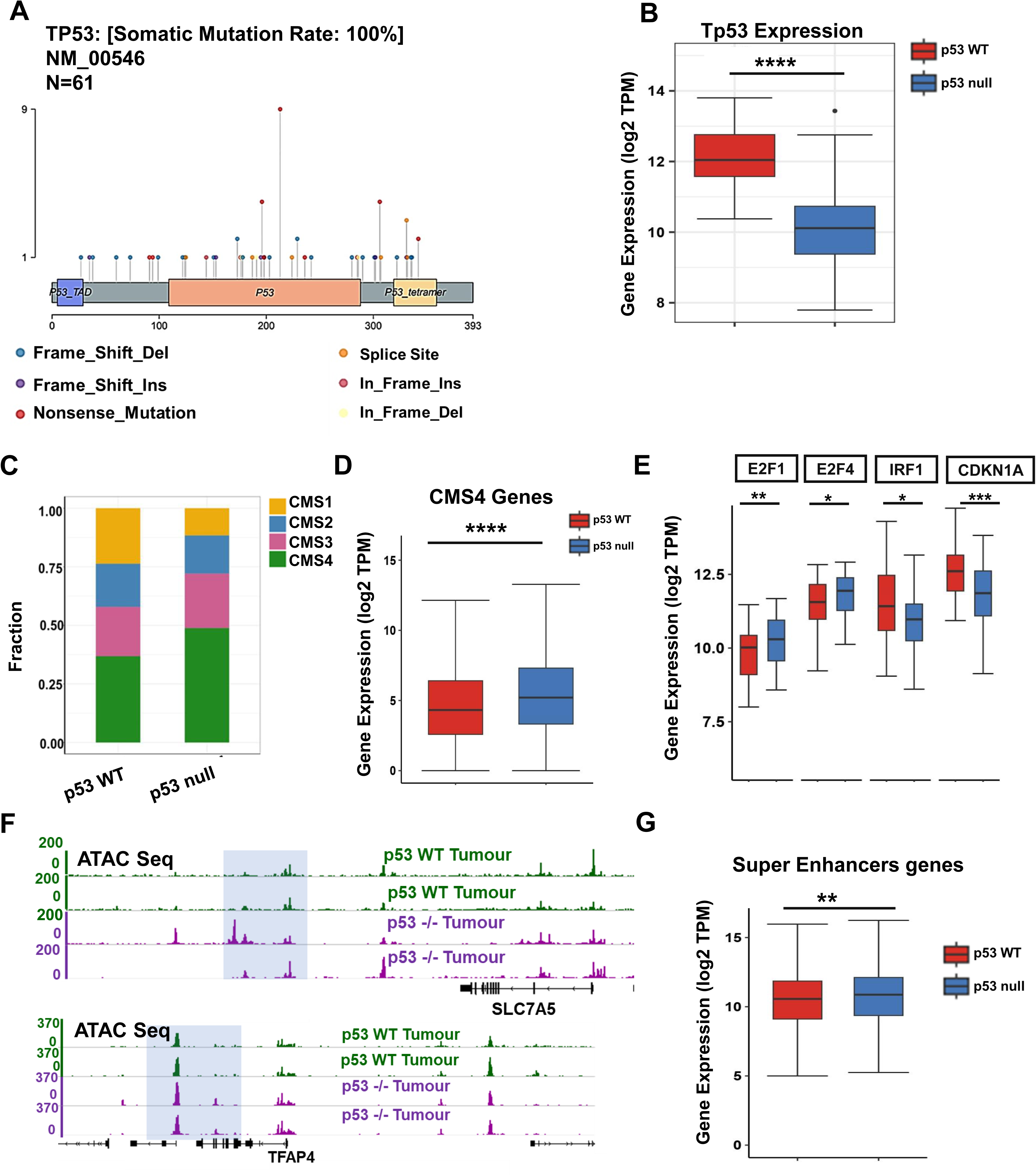
Validation of the identified epigenetic regulation in p53 null tumors derived through TCGA-COAD **A.** Lollipop plot represents different sites of p53 mutation in p53 null tumours through COAD cohort (N=61). TP53 deleterious tumours includes deletion, insertions and nonsense mutations. **B.** Box plot representing gene expression profile of Tp53 in p53WT patients and p53 null COAD patients (t-test was performed for the statistical analysis, p value> 0.05, ns; p value<=0.05, *; p value<=0.01, **; p value<=0.001, **). Patients harbouring deleterious mutation of Tp53 show a decrease in the expression profile of Tp53 at transcript level. **C.** Stacked bar plot representing CMS classification in WT p53 and p53null COAD patients based on CMS classifier. Patients with p53 null mutation shows higher fraction of tumour population enriched with CMS4(green). **D.** Boxplot representing the gene expression profile of CMS4 signatures in p53 WT (red) and p53null patients (blue) (t-test was performed for the statistical analysis, pvalue> 0.05, ns; p value<=0.05,*; p value<=0.01,**; p value<=0.001,***). p53 null tumours are associated with significantly higher expression of CMS4 gene signatures as compared to p53 WT tumours. **E.** Box plot representing gene expression profile of E2F1, E2F4, CDKN1A and IRF1 in p53WT tumours (red) and p53 null tumours (blue) (t-test was performed for the statistical analysis, pvalue> 0.05, ns; p value<=0.05, *; p value<=0.01, **; p value<=0.001, **). Tumours with p53 null tumours are associated with significant higher expression of E2F1, E2F4 and significantly lower expression of CDKN1A and IRF1. **F.** IGV screenshot representing ATAC signals in p53 WT (green) and p53 null (purple) tumours around the super enhancers of oncogenes SLCA7A5 and TFAP4. Oncogenic super-enhancer regions of SLCA7A5 and TFAP4 correlates with increased accessibility in p53 null tumours. **G.** Boxplot representing the gene expression profile of the identified gained super enhancer genes in p53WT (red) and p53 null tumours(blue) (t-test was performed for the statistical analysis, pvalue> 0.05, ns; p value<=0.05, *; p value<=0.01, **; p value<=0.001,**). A significant increase in the expression profile of oncogenic superenhancer genes in p53 null tumours was observed compared to p53WT tumours.

Conventionally, colorectal cancers have been classified into 4 different consensus molecular subtypes based on both genomic and epigenomic features, CMS1, CMS2, CMS3 and CMS4 (Guinney et al., 2015). We attempted to classify both our WT and p53 null cohort derived from TCGA-COAD into different CMS subtype using two known available tools/packages CMS classifier ((Guinney et al., 2015)) and CMS caller (Eide et al., 2017). We observed that a large fraction of p53 null mutation CRC patients belonged to CMS4, while the most of the p53 WT tumours belonged to CMS1 subtype (Fig 7C,Fig S6 A,B). CMS4 tumour subtype is known to be associated with epithelial mesenchymal transition (EMT) and gene signatures associated with the activation of transforming growth factor β (TGF β) signaling (Guinney et al., 2015). This suggests that p53 null tumours have more metastatic potential and correlates with poor survival as compared to CRC patients with p53 WT tumours. We also compared the expression of these CMS4 gene signatures in p53 WT and p53 null tumours, and observed that the CMS4 gene signatures show higher expression in p53null tumours as compared to p53 WT tumors (Fig7D). This again suggests that the expression patterns in p53 null CRC patients correlate more with higher metastasis and oncogenesis as compared to p53 WT CRC patients, which also might explain the observed rates of their lower survivability. In order to further explore the changes in expression profiles of the transcriptional circuitry observed with the loss of p53 in CRC patients, we compared the expression profiles of various E2Fs, IRF and CDKN1A involved in p53 signalling in p53WT and p53null tumours from the TCGA-COAD cohort. Interestingly, we observed that the expression profiles of E2Fs were significantly upregulated including E2F1 and E2F4 and CDKN1A was found to be downregulated (Fig7E), suggesting that loss of p53 upregulates various E2Fs mostly through an indirect mechanism involving CDKN1A.

We also utilised the ATAC sequencing data available in the TCGA-COAD to determine the accessibility around the gained and lost super-enhancer regions in p53 WT(N=2) and p53 null CRC tumours(N=2). We observed that the gained super-enhancer regions correlate with increased accessibility while the lost super-enhancer regions correlate with decreased accessibility in p53 null tumours as compared to p53 WT tumours (Fig 7F, Fig S6D, S6E). In order to understand the oncogenic gained super enhancer associated gene signatures across p53 null and p53 WT CRC patients, we analysed the overall expression profile of the 117 oncogenes associated with gained super enhancers in patients with p53WT tumours and p53 null tumours and found that these oncogenic signatures were significantly upregulated in p53 null CRC patients as compared to p53 WT CRC patients (Fig7G).

Furthermore, comparison of the TCGA-COAD patients tumours displaying high (upper quartile) expression of the oncogenic signature associated with gained super enhancers Vs those with low (bottom quartile) expression revealed that high oncogenic signature expression of 101 genes out of 117 were significantly associated with lower survivability (Fig S6F). We next performed ssGSEA on the differentially expressed oncogenes in p53 null and p53 WT tumours and found that these genes are involved in MYC activation pathways (Fig S6E) in p53 null tumours as compared to p53WT tumours implying that low p53 expression activates MYC pathway in colon tumours, which are mostly regulated by distal regulatory super enhancer regions. The epigenetic regulation observed in colorectal cancer cells through the enhancer circuitry upon loss of p53 highly correlates with our observations on colon cancer patients too.

## Discussion

A multistep genetic model by Vogelstein suggesting accumulation of several genetic and epigenetic alterations in key genes involved in silencing of TSGs and activation of oncogenes (Fearon & Vogelstein, 1990) has established the first understanding of tumour suppressor function in colorectal cancer. The loss or gain of Function (GoF) mutation in p53 is a late event in cancer progression that drives the cancer from adenomatous stage to carcinoma stage (Schwitalla et al., 2013, Onuma et al., 2013).

Pioneering studies investigating the chromatin pathways aligning with genomic changes in p53 mutated breast cancer cells have shown that the genome wide binding of GoF p53 shows preferential binding to the MLL1 and MLL2 genes which are a part of the COMPASS complex and are functionally related to histone methylation (H3K4me3). Inhibition of these enzymes using Menin (COMPASS inhibitor) resulted in the reduction of tumour population validating these epigenomic signatures (Zhu et al., 2015). Such studies have strongly emphasized a cross talk between the epigenomic circuitry with genetic events orchestrating cancer progression.

Though alterations in regulatory landscape of the cancer genome have been shown to play a key role in metastasis, reprogramming of these regulatory circuits and the specific roles of distant transcriptional regulators upon loss of p53 in oncogenesis and metastasis have not been well established. In this work, we have extensively characterized the alterations in the enhancer landscape, oncogenic super enhancers and the transcriptional circuitry leading to metastasis upon loss of p53 in colorectal cancer.

Through an extensive experimental and computational approach involving ChIP and RNA sequencing, we have shown a global increase in H3K27ac mark upon p53 loss distributed mostly around the promoter and distal regions of the colorectal cancer genome. This observation corroborates with increase in chromatin accessibility at both the promoters and enhancers upon EMT induction using TGFβ treatment in primary mammary epithelial cells (NMuMG cells) (Timmers et al., 2007) implying that a genome wide increase in H3K27ac at both the promoters and enhancers may be responsible for metastatic progression in colorectal cancer upon p53 loss.

On functionally integrating these genes associated with gained and lost H3K27ac sites at promoters and enhancers through overlap of the ChIP and RNA sequencing data, we observed an upregulation of E2F and MYC targets at the gained enhancer sites and a down regulation of interferon signalling at the lost enhancer sites.

Our analysis of the differential motifs regulating these signatures revealed that gained H3K27ac sites were enriched with more GC rich motifs with distinct transcriptional factors involved with metastasis and stemness including E2Fs, SP1, MAZ, PATZ1 while the lost H3K27ac sites had more AT rich motifs with transcriptional factors like GATA2, IRFs and P53 responsible for restoring epithelial identity. Functional integration of the identified transcription factors at these distinct regions with the transcriptome sequencing reveals an upregulation of E2Fs, MAZ and PATZ1 suggesting a novel regulatory circuit mediated by active enhancer mark upon p53 loss in colorectal cancer.

Interestingly, we observed that the upregulation of the identified transcriptional factors including E2F1, E2F2, MAZ and PATZ1 at the enriched regions correlates with elevated levels of H3K27ac marks around their promoter regions. We therefore hypothesize that loss of p53 promotes cancer metastasis and stemness by remodelling the levels of the active enhancer mark H3K27ac around the promoters and enhancers of various E2Fs like transcriptional factors which in turn have positive feedback loop upon loss of p53.

This model corroborates with the finding where E2F mediated target expression and cellular proliferation have been reported to be regulated by a WT p53 dependent negative feedback loop (Timmers et al., 2007). It is known that both E2F-1 and p53 utilize p300 as a transcriptional co- activator, and the competitive interaction of p53 or E2F-1 for p300 determines functional outcome of cellular proliferation (Lee et al., 1998). We observed an elevation of H3K27ac around the promoters and enhancers of various E2F targets correlating with their high expression upon p53 loss. We therefore hypothesize that upon loss of p53, E2F-1 might utilize p300 and in turn have a positive feedback regulation and thus promotes E2F mediated cancer stemness and metastasis.

We also establish an indirect upregulation of histone methyl transferases EZH2 and SuV39H1 upon loss of p53. We establish that the loss of p53 indirectly promotes metastasis by E2F and MAZ driven transcriptional circuitry around the gained H3K27ac sites. The E2F driven transcriptional circuitry include histone methyl transferases EZH2 and SuV39H1, which serve as oncogenes in promoting colorectal cancer. Our data also demonstrates an increase in expression of p73 which might be due to increase in E2F1 binding around the promoters associated with increased H3K27ac levels, suggesting an indirect gain of active enhancers upon loss of p53. The loss in levels of H3K27ac at the distal regions of IRF coinciding with reduced interferon gamma responses in the CRC genome suggests that the transcriptional circuitry driven by WTp53 also involve various other tumour suppressive genes including IRFs.

Super enhancer formation is a distinct characteristic of cancer genomes during progression and metastasis. Interestingly, our experiments reveal that distal super enhancer regions show an overall gain of super enhancer regions upon p53 loss correlating with increased expression of various oncogenes. The presence of Zinc Finger Nucleases - like transcriptional regulators at the gained super-enhancer regions including ZNF281, E2F and KLFs, suggest an indirect role of p53 in formation of these gained super-enhancer regions. The observed increase in spreading of H3K27ac around these gained super-enhancer regions upon loss of p53 as evidenced by increase in peak width and also on the number of enhancer peaks per super-enhancers highlights a possible metastatic progression upon loss of p53 accompanied by increase spreading of these typical enhancers.

The gained oncogenes associated with super enhancers further led us to investigate the overall expression of tumour suppressor genes and oncogenes. We observed that loss of p53 accompanies decrease in tumour suppressor expression correlating with gained oncogenic expression and the levels of H3K27ac. An in-depth analysis of the motifs around the promoters of oncogenes and tumour suppressors, accompanied by gained and lost H3K27ac sites further establishes that wildtype p53 functions as direct activator for various TSGs while the oncogene activation is mainly driven through E2F/SP1 like transcription factors, highlighting a direct activating role of WTp53 on tumour suppressor genes while an indirect repressive role on oncogenes.

In order to validate and understand the epigenetic circuitry in colon tumours, we identified 61 tumours with p53 null mutation and 61 WTp53 colon tumours. Initial consenus molecular subtyping (CMS) of colon tumours has revealed enrichment of p53 mutations in CMS2 followed by CMS4 subtype, however stratification of tumors based on distinct p53 mutation subtypes have not been explored (Guinney et al., 2015). This work for the first time has identified p53 null tumours to be enriched in CMS4 subtype, correlating with increase in the expression profile of known CMS4 genes. Furthermore, recent epigenomic classification with H3K27ac in colon cancer has revealed that the EpiC2 subtype which overlaps with CMS4 subtype shows higher enrichment of H3K27ac as compared to the other EpiC clusters (Orouji et al., 2022) which suggest that p53 null tumors in colon cancer are accompanied by increased H3K27ac levels. Our RNA sequencing analysis of these patients further identified significant upregulation of E2F1 and E2F4 in p53 null tumours, highly correlating with signatures observed in p53-/- CRC cells. We also observed an upregulation of super-enhancer associated genes in p53 null tumours and gained accessibility at the gained super-enhancer regions with lost accessibility in lost super enhancer regions in p53 null tumours.

In summary, this work identifies a hitherto unknown novel regulatory mechanism mediated by active enhancer H3K27ac mark upon loss of p53. We show that the loss of p53 induces gain in active enhancer mark at promoters and distal intergenic regions which is mostly due to an indirect effect of p53. We have identified a distinct E2F mediated regulation of cancer stemness and metastasis effected by reprogramming of H3K27ac with loss of p53. We have also found an indirect effect of p53 in upregulation of known E2F targets EZH2 and SuV39H1, the methyl transferases for H3K27 and H3K9 methylation. Interestingly, we have also observed a novel gain in oncogenic super-enhancers upon loss of p53 in colorectal cancer cells which observation has also been validated in patient tumors harbouring p53 null mutation.

Our data highlights that the combination of p53 loss and increase in active H3K27ac regulation activates oncogenic pathways including MYC and E2F targets, suggesting an epigenetic reprogramming upon loss of p53 in CRC. Our data clearly establish a powerful interplay between the epigenetic machinery and the tumour suppressor network in modulating cancer progression through enhancer regulation. This provides a clinical framework of devising enhancer based epigenetic therapy for patients with p53 deletions. Synergistic therapeutic strategy with EZH2, SuV39H1 and BRD could be a promising strategy in reprogramming of CMS population through alteration in H3K27ac levels and the change in tumour microenvironment in patients harbouring p53 loss/mutation. This also provides a possibility of modulating super enhancer function to overcome drug resistance over chemotherapeutic regimes in colorectal cancer.

## Materials and Methods

### Cell Culture

Colon adenocarcinoma cell lines with wild type p53 (HCT116 p53WT) procured from NCCS, Pune and colon adenocarcinoma cells with deletion of p53 (HCT116 p53-/-) (Kind gift from Bert Vogelstein Lab, Johns Hopkins University and Jayant Sarkar Lab, CDRI, Lucknow) were maintained in Dulbecco’s Modified Eagle’s Medium and Hams-F12 (DMEM-F12) (#12500062 Invitrogen-Gibco) supplemented with 10% FBS (#16000044 Invitrogen-Gibco) and 1% penicillin-streptomycin (#1507070063 Invitrogen-Gibco). The cells were maintained in humidified environment at 37°C in presence of 5% CO2.

### Chromatin immuno Precipitation assay and library preparation

2.5 million cells were seeded and cultured to attain 80-90% confluency in 100mm dishes. The cells were crosslinked with 1% formaldehyde (#219404790 MP BIO) at room temperature for 10 mins with constant shaking. Glycine was added to a final concentration of 125 mM to quench the formaldehyde for 5 mins. Cells were washed thrice with 1X ice cold PBS. Cells were scraped in 3 ml of 1X PBS and pelleted down at 2K rpm for 5 mins at 4°C and kept at -80° C. Nuclear lysis buffer (L2) (50 mM Tris-HCl pH 7.4, 1% SDS, 10 mM EDTA pH 8.0) supplemented with 1X PIC was added to the nuclear pellet and incubated on ice for 10 mins. Samples were sonicated using Covaris-S220. The cell lysate was cleared by centrifuging samples at 12K rpm for 12 mins. 100 ug of sheared chromatin was taken for each IP. Sheared chromatin was diluted by adding Dilution Buffer (DB) (20 mM Tris-HCl pH 7.4, 100 mM NaCl, 2 mM EDTA pH 8.0, 0.5% Triton X-100) supplemented with 1X PIC in 1:1.5 (1 volume of sheared chromatin and 1.5 volumes of Dilution Buffer) ratio. 10% of diluted chromatin was set aside as Input. 2µl of antibody(H3K27ac(#8173S)) was used for each IP. Dynabeads (Invitrogen-#140004D) were prepared by blocking in 1% BSA prepared in 1X PBS at 4°C for 1 hour followed by washing with 1X PBS. Immunoprecipitated DNA was collected by adding 14µl of BSA blocked beads to each sample and incubated at 4°C for 4 hours. Beads were collected, flow through was discarded and 800 µl of Wash Buffer I (20 mM Tris-HCl pH 7.4, 150 mM NaCl, 0.1% SDS, 2 mM EDTA pH 8.0, 1% Triton X-100) was added. Washings were carried out at 4°C on a rocking platform. Washings were sequentially repeated with Wash Buffer II (20 mM Tris-HCl pH 7.4, 500 mM NaCl, 2 mM EDTA pH 8.0, 1% Triton X-100), Wash Buffer III (10 mM Tris-HCl pH 7.4, 250 mM LiCl, 1% NP-40, 1% sodium deoxycholate, 1 mM EDTA pH 8.0) and 1X TE (10 mM Tris pH 8.0, 1mM EDTA pH 8.0). Immunoprecipitated DNA was eluted by adding 200 μl of Elution Buffer (100 mM NaHCO3, 1% SDS) for 45 mins at 37°C in a thermomixer with rpm of 1200. Eluate was transferred to fresh tubes and 14 µl of 5M NaCl was added and kept overnight at 65°C. Immunoprecipitated DNA was purified by Phenol:chloroform:isoamyl alcohol (Ambion-AM9732) method, followed by ethanol precipitation. The final dried DNA pellet was dissolved in 15µl of 1X TE. 1ng of the DNA was used for preparation of ChIP libraries using NEBNext® ltra™ II DNA Library Prep with Sample Purification Beads (#E7103L) and subjected to sequencing using illumina Hi Seq 2500 sequencing platform, 1×55 bp sequencing read length.

### RNA-Seq library preparation and sequencing

50,000 cells were cultured as described above for isolation of RNA for sequencing. Cells were lysed with 500µl of RNA iso reagent (#9108 Takara). 1/5^th^ volume of chloroform was added to samples, briefly vortexed and centrifuged at 12K rpm for 12 mins at 4 degrees. The aqueous phase was carefully collected and transferred to fresh tubes. 0.7 volumes of isopropanol were added to samples and incubated at room temperature for 10 mins to precipitate the RNA. The samples were centrifuged at 12K rpm for 12 mins, supernatant was discarded without disturbing the pellet. The pellet obtained was washed with 80 % ethanol. Pellet was allowed to dry for 20 mins and dissolved in DEPC treated water. 1ug of RNA was used for library preparation. RNA sequencing libraries were prepared with Illumina-compatible NEBNext® ultra™ II Directional RNA Library Prep Kit (New England BioLabs). High throughput sequencing was performed on Illumina Hi Seq2500 platform.

#### ChIP-Seq data processing and analysis

Fast QC was done for all the samples to check for adapter content, PHRED score and GC content. Adapter content was removed using Trimmomatics (v 0.39). Trimmed reads were later mapped to the human genome (hg38) using Bowtie2(v 0.2.4.1) (Langmead & Salzberg, 2012). Sam files were converted into sorted and indexed bam files using samtools. Peak calling was performed using MACS2 (Zhang et al., 2008) with a p-value threshold of 5E-10 and --SPMR normalization. For each condition, the corresponding input samples were used for background normalization. The reproducibility between the replicates and consensus set of peaks was derived using irreproducibility discovery rate (IDR) (Q. Li et al., 2011) (ver 2.0.4.2) analysis using the recommended parameters. To compare the ChIP sequencing data sets of the H3K27ac enhancer mark across different conditions, the consensus set of peaks obtained in each condition were merged using the “merge” function from bed tools ((Quinlan & Hall, 2010)). The density of reads in each merged region was quantified using normalized signal per million reads. The gained and Lost sites were defined on the basis of normalized signal per million reads of H3K27ac for which fold change (FC) was larger than >1. The boxplot comparing the H3K27ac enrichment in different conditions was drawn by R-package ggplot2 and quantified for significance using normalized CPM counts by Wilcoxon test. Bigwig files were created using Deep tools bamcoverage (Ramírez et al., 2014). The heat map was drawn by plotHeatmap. Profiles were obtained on a region of ±2 kb from the center of peaks, and the average scores were plotted to generate averaged read density around peaks using Deeptools plotProfile (Ramírez et al., 2014). The representative genome browser snapshot was shown by IGV.

### Identification of enhancers

The ChIP sequencing peaks corresponding to H3K27ac within 2 kb of transcription start sites were labelled as active promoter peaks while those peaks beyond +/- 2kb were labeled as enhancer peaks. A consensus set of enhancer regions in each condition was derived using IDR (v 2.0.4.2) ((Q. Li et al., 2011)). To identify differentially enriched enhancers, a consensus set of enhancers obtained in each condition were merged using the “merge” function from bedtools (Quinlan and Hall 2010). The density of reads in each merged region was quantified using normalized signal per million reads. The gained and lost enhancers were defined on the basis of normalized signal per million reads of H3K27ac for which fold change (FC) was larger than >1. The normalized CPM H3K27ac signals were then quantified around gained and lost enhancers using ‘R’. The boxplot comparing the H3K27ac enrichment between enhancer peaks in different conditions was drawn by R-package ggplot2 and the p-value was calculated by Wilcoxon test.

### Identification of super-enhancers

To identify super-enhancers (SE), the program ‘‘findPeaks’’ in the HOMER version 4.10.3 ((Heinz et al., 2010)) was used with the option ‘‘-style super” parameters for each replicate under each condition. Style super merges identified peaks found within 12.5 kb into continuous regions. The “enhancer score” for each region is defined by the number of experimental reads minus the input reads normalized for sequencing depth. All regions were then sorted by their score and plotted by their relative rank and score (0-1). Enhancer regions beyond which the slope of the line reaches 1 are considered SEs. The elbow of the signal curve was determined as the slope = 1 and super-enhancers were classified based on a criterion that the signal was greater than the slope (1) respectively (Whyte et al., 2013). Signal intensity of the determined peaks was calculated and normalized into CPM. To identify differentially enriched super-enhancers, all SEs identified in either p53WT and p53-/- CRC cells were merged using ‘‘mergeBed’’ (bedtools version 2.26.0 package). “Bedmap” in the bedtools was later used to quantify normalized signal per million reads across the merged super-enhancer regions in each replicate. SEs within log2FC >=0.5 or log2FC<=-0.5 signals were classified as Gained and Lost super-enhancers respectively.

### Gene assignment to enhancers and super enhancers

The identified gained and lost enhancers and super enhancers regions were associated with genes within 1Mb distance using the Genomic Regions Enrichment of Annotations Tool (GREAT; http://great.stanford.edu/public/html/index.php) with default settings. The active genes with log2FC >= 0.5 in gained H3K27ac signals were defined as genes that gained the signal/upregulated genes associated with enhancers and super-enhancers. Similarly, the genes with log2FC <= -0.5 in lost H3K27ac signals were classified as signal-lost genes. Boxplots representing distribution of gene expression profile (FPKM) values for gained and lost enhancers and super-enhancers were plotted using ggplot2 in R. The statistical analysis was done using a paired ‘t’ test.

### Motif Analysis

The differential motif enrichment analysis was performed for the following comparisons: (1) gained H3K27ac sites versus lost H3K27ac sites and *vice-versa*, (2) promoters of oncogenes associated with gained H3K27ac sites versus promoters of tumor suppressor genes associated with lost H3K27ac sites and *vice versa* (3) Gained super-enhancers Vs lost super-enhancers and *vice versa*. The XSTREME package (Bailey et al., 2015) was used to identify motifs. By default, XSTREME reports 6- to 15-mer motifs whose E-value ≤ 0.05. The program uses Fisher’s exact test or the binomial test to determine the significance of each motif found.

### RNA-Seq data processing and analysis

The paired end RNA seq reads were mapped to GRCh38/hg38 reference genome using STAR (v2.5.3a) aligner (Dobin et al., 2013). To perform differential expression analysis, raw read counts matrix was estimated by HTseq count. Differential expression analysis was carried out using R/Bioconductor package DESeq2(v1.22.1) (Love et al., 2014). Genes with p value less than 0.05 and absolute fold change more than |0.5| were considered as differentially expressed genes. Heat maps were generated using z- scores with the R package pheatmap (v1.0.10). Functional annotation based on gene ontology was performed using Enrichr.

### Gene Set Enrichment Analysis (GSEA)

We have performed GSEAPreRanked (subset of GSEA) (Subramanian et al., 2005a) of the genes ranked on the basis of gene expression values (higher to lower) and the significance to identify the significantly enriched biological processes and pathways associated with a set of differentially expressed genes. The gene sets used in the analysis were obtained from the set of differentially expressed genes. The enrichment analysis was performed using the default settings and statistically significant enrichment results were determined based on the false discovery rate (FDR) adjusted p-value threshold of 0.05.

### Identification of p53null and p53 mutant CRC patients from TCGA and gene expression analysis

The mutations harboured by the cohorts in the TCGA -COAD project was classified into p53 null mutation(N=61), and p53 WT(N=61) based on the somatic mutations reported for colon cancer (TCGA-COAD). The p53 null group was completely devoid of missense mutations and contained frame-shift, nonsense and splice-site mutations, which occur as a consequence of intragenic nucleotide insertions or deletions and/or single base-pair substitutions, which are expected to affect protein encoding reading frames and hence associated with unstable transcripts. The raw expression counts of protein-coding genes for colon cancer samples were obtained from the GDC portal using TCGAbiolinks. Further, the count matrix was normalized (using TCGAanalyze_Normalization function) and filtered out genes with low expression count (using TCGAanalyze_Filtering with method = quantile, and qnt.cut = 0.25). The differential expression analysis was carried out using limma method through TCGAanalyze_DEA function (with method = glmLRT). Genes with |log2foldchange| > 0.5 and FDR (q-value) < 0.1 were considered significant and differentially expressed. The difference in expression in the TCGA samples was computed using ‘t’ test. Single sample gene set enrichment analysis was performed on the differentially expressed genes using GSEA tool ((Subramanian et al., 2005).

### CMS classification

Ensembl IDs were translated to Entrez IDs with the biomaRt Bioconductor package. In order to achieve accuracy in classification, the random forest CMS classifier (Guinney et al., 2015) and CMS caller (Eide et al., 2017) were applied individually on the p53 WT patients and p53null patients (TCGA-COAD). Gene expression matrices on log2-scale were normalized to z-scores classified with the nearest template prediction approach in the R package CMScaller with an FDR-threshold <0.05 (v.2.0.1). CMS class was assigned according to the default settings (minCor = 0.15, minDelta = 0.06) in CMS caller. To obtain the original CMS labels for TCGA-COAD samples, the random forest CMS classifier was also applied to the whole TCGA-COAD dataset. To obtain CMS4 based gene signatures in TCGA-COAD we used differential TCGAanalyze_DEA function (with method = glmLRT) and performed differential gene expression analysis (CMS4 cohort vs others). Box plots representing gene expression profile in p53WT patients Vs p53 null patients was plotted using ggplot2 in R. Student ‘t’ test was used to evaluate the significance in expression profiles in the two cohorts (p53WT COAD patients Vs p53null COAD patients).

### TCGA-ATAC Seq Analysis

TCGA-COAD type-specific bigwig files and raw counts of ATAC-Seq peaks (called at the individual sample-level) were obtained from Corces & Granja et al (Corces et al., 2018). (https://gdc.cancer.gov/about-data/publications/ATACseq-AWG). 2 p53null tumors and 2 p53WT colon tumours were identified and their normalized bigwig files were used to quantify accessibility around the gained and lost super-enhancer regions.

### Survival Analysis for TCGA COAD patients

To investigate the clinical significance of the oncogenes associated with gained super-enhancers in the p53-/- CRC cell line, survival analysis was performed on the TCGA-COAD (The Cancer Genome Atlas - Colon Adenocarcinoma) cohort for 270 colorectal cancer patients. The Kaplan-Meier curves were generated using GEPIA2 to screen the hazard ratio of genes which were selected with p-value<=0.1 and the “median” as the group cutoff. All the statistical analysis was performed in R4.4.1.

## Supporting information

All_Supplementary

## Acknowledgement

- The authors acknowledge the sequencing facility at C-CAMP, BLiSc, Bangalore.
- Facility at Notani lab at NCBS-TIFR for ChIP sequencing

## Data Availability

ChIP sequencing data generated in this study are submitted in GEO database under BioProject PRJNA1227440. RNA sequencing data generated in this study are submitted under BioProject PRJNA1016399.

