## Supplementary material for "Loss of tumour suppressor p53 rewires enhancer landscape and governs oncogenic progression": All_Supplementary

Fig S1

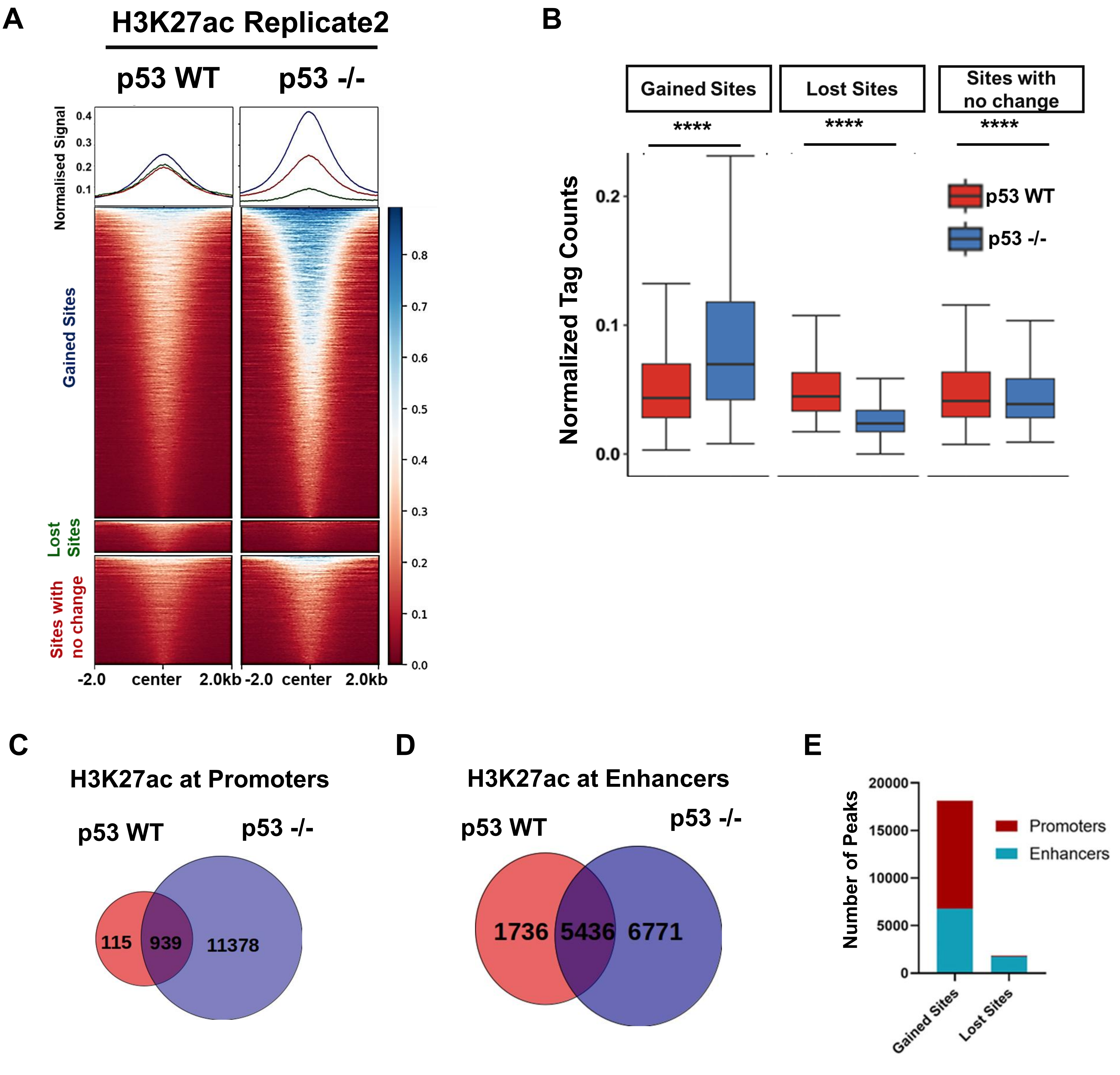
A
H3K27ac Replicate2
p53 WT
p53 -/-
Normalised Signal
Gained Sites
Lost Sites
Sites with no change
-2.0 center 2.0kb -2.0 center 2.0kb
B
Gained Sites
Lost Sites
Sites with no change
****
Normalized Tag Counts
p53 WT
p53 -/-
C
H3K27ac at Promoters
p53 WT
p53 -/-
115
939
11378
D
H3K27ac at Enhancers
p53 WT
p53 -/-
1736
5436
6771
E
Number of Peaks
Promoters
Enhancers
Gained Sites
Lost Sites

Fig S2

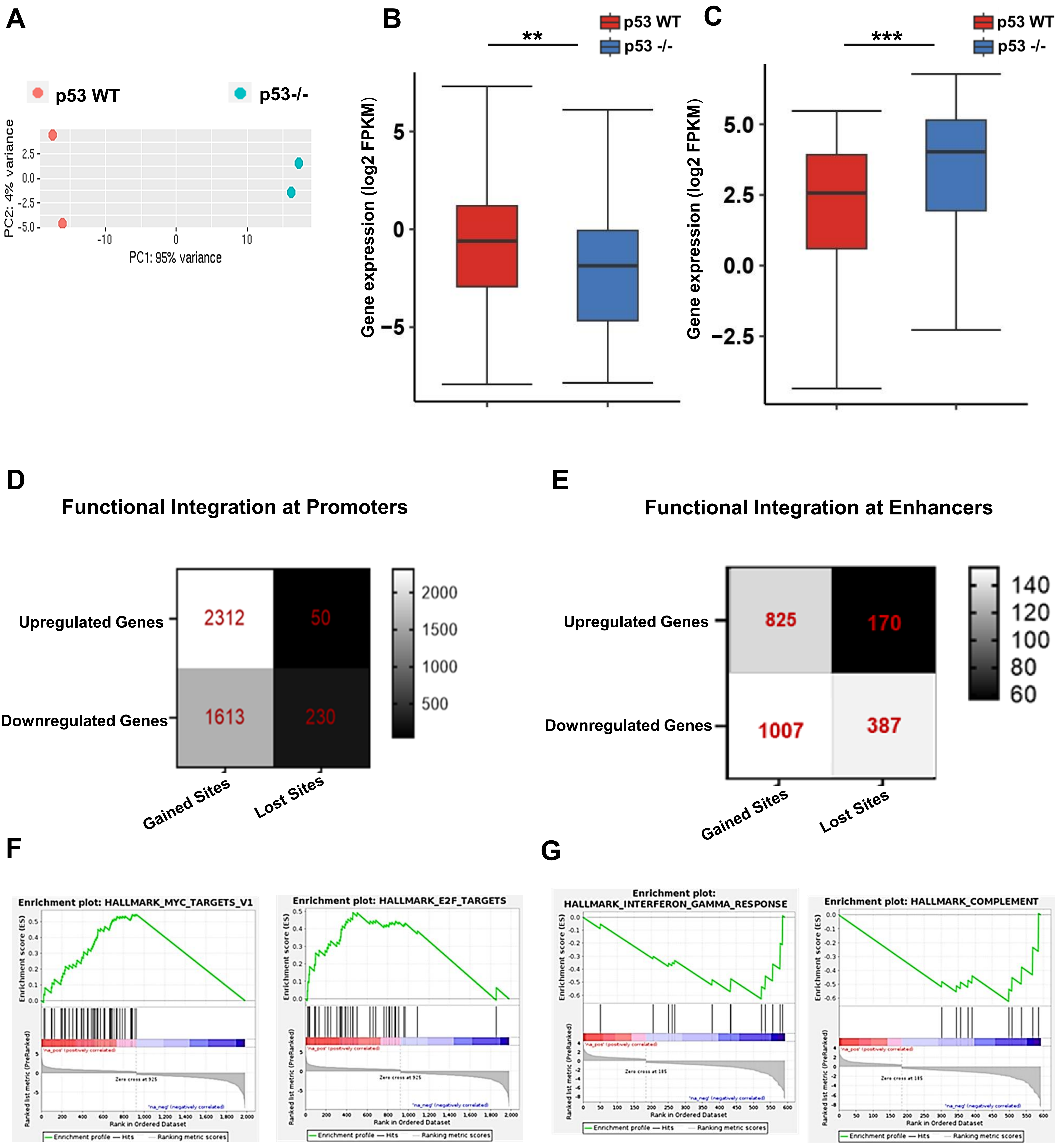
A
p53 WT
p53-/-
PC2: 4% variance
PC1: 95% variance
B
p53 Signalling
**
p53 WT
p53 -/-
Gene expression (log2 FPKM)
C
MYC Targets
***
p53 WT
p53 -/-
Gene expression (log2 FPKM)
D
Functional Integration at Promoters
Upregulated Genes
Downregulated Genes
2312
50
1613
230
Gained Sites
Lost Sites
E
Functional Integration at Enhancers
Upregulated Genes
Downregulated Genes
825
170
1007
387
Gained Sites
Lost Sites
F
Enrichment plot: HALLMARK_MYC_TARGETS_V1
Enrichment plot: HALLMARK_E2F_TARGETS
G
Enrichment plot: HALLMARK_INTERFERON_GAMMA_RESPONSE
Enrichment plot: HALLMARK_COMPLEMENT
Enrichment score (ES)
Ranked list metric (PreRanked)
Rank in Ordered Dataset

**Fig S3**

**A**

| TF | Consensus | TP% | p value |
|---|---|---|---|
| SP2 | | 60.39 | 1.76e-354 |
| AP2B | | 68.98 | 1.88e-332 |
| E2F5 | | 73.2 | 7.49e-331 |
| SP1 | | 51.76 | 5.88e-324 |
| E2F4 | | 62.85 | 5.42e-304 |
| E2F1 | | 66.18 | 2.72e-292 |
| ZN341 | | 52.79 | 3.11e-324 |

**B**

| TF | Consensus | TP% | p value |
|---|---|---|---|
| ZN394 | | 82.65 | 1.03e-243 |
| GATA2 | | 77.91 | 6.94e-221 |
| ANDR | | 77.73 | 4.43e-215 |
| NKX6-1 | | 80.13 | 5.92e-215 |
| PIT1 | | 75.21 | 4.61e-211 |
| MEF2A | | 74.49 | 3.18e-206 |
| MEF2D | | 75.09 | 5.9e-204 |

**C**

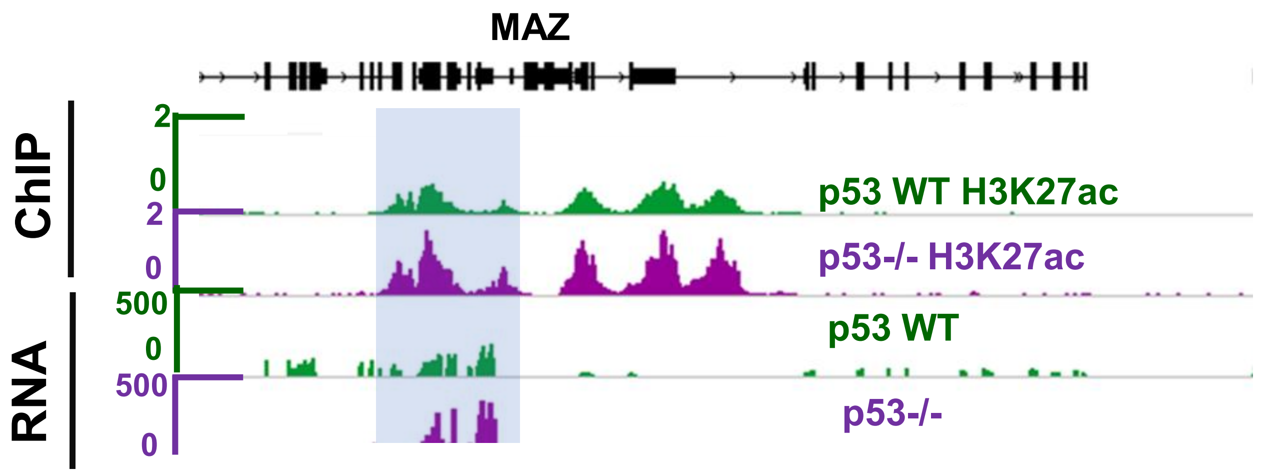


**D**

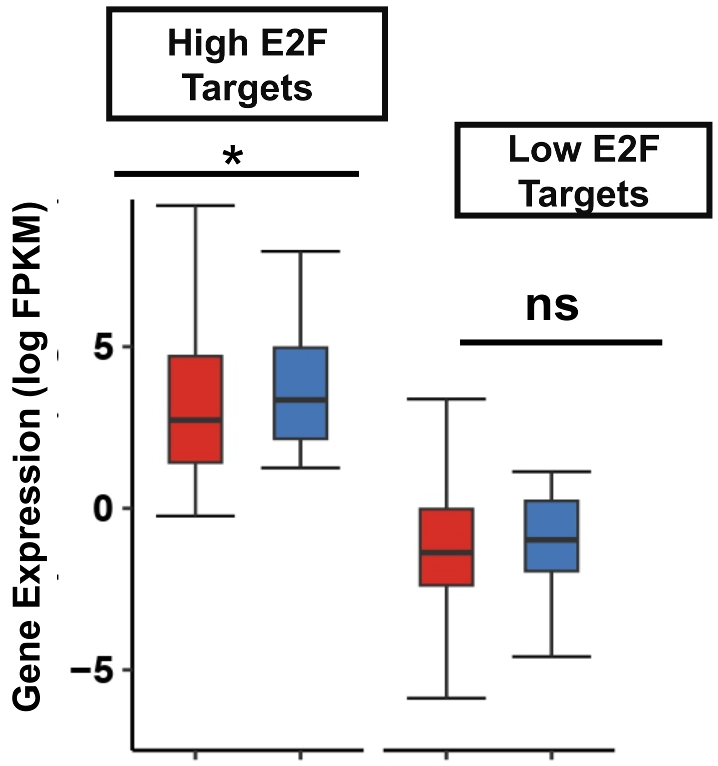


**E**

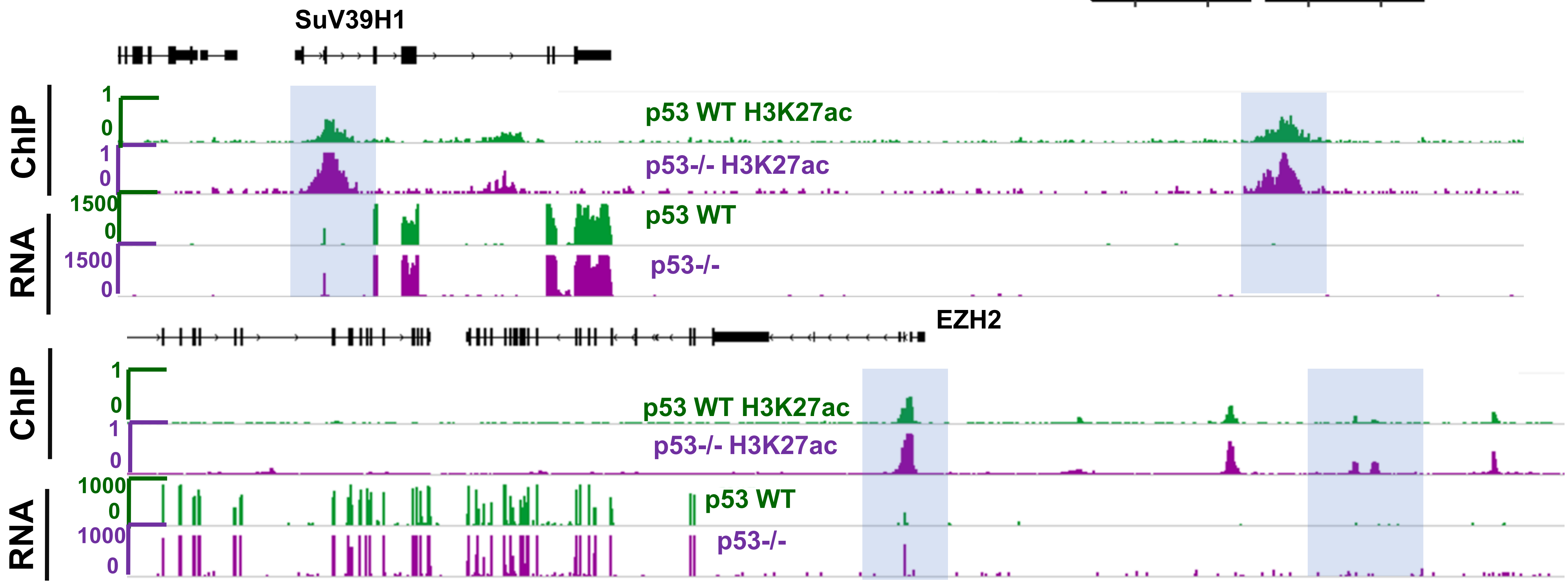

### Fig S4

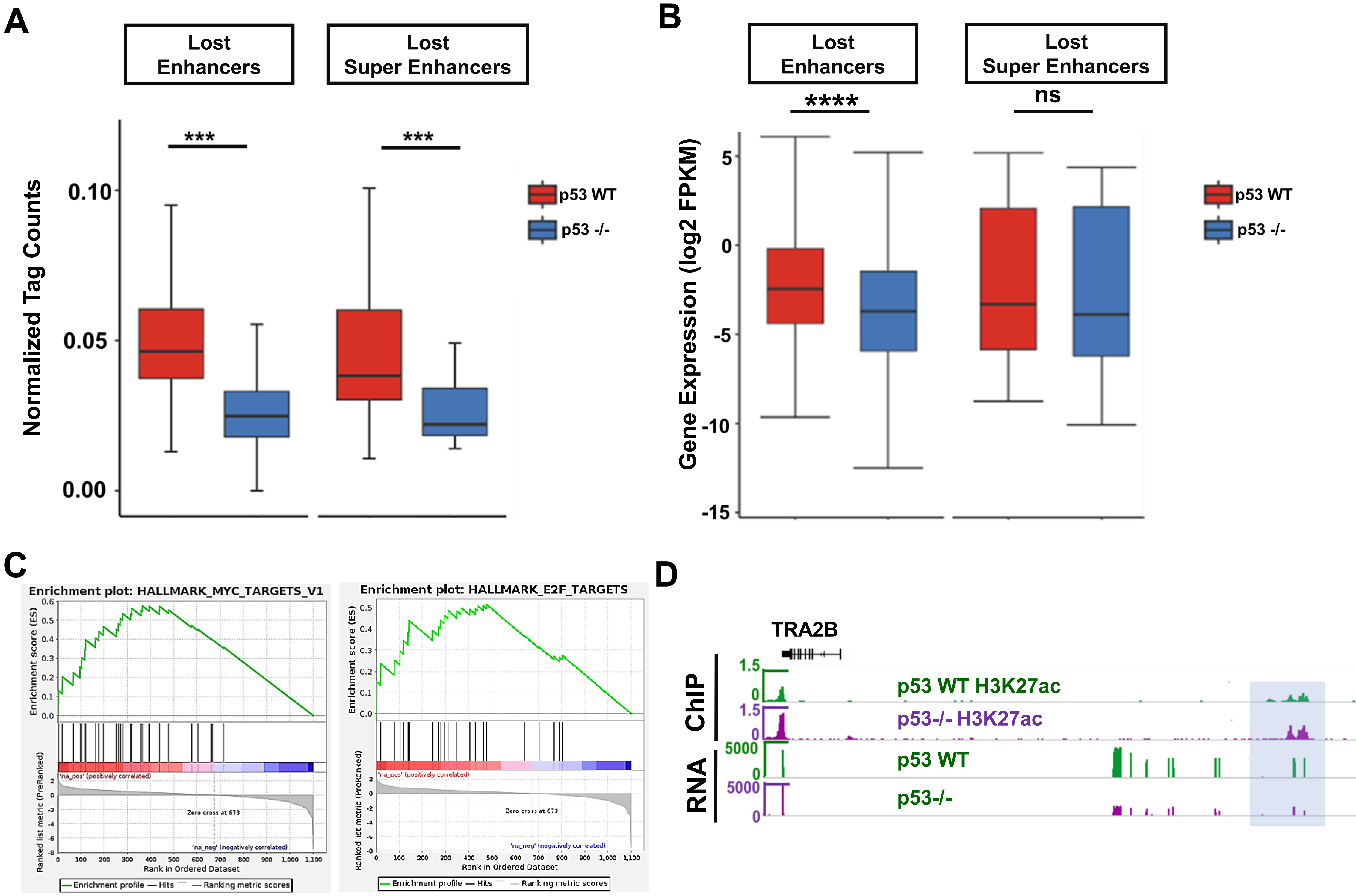


**E**

#### Gained Super Enhancers

| TF | Consensus | TP% | p value |
|---|---|---|---|
| ZN281 | 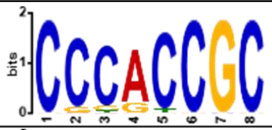 | 91.2 | 7.45e-5 |
| ZBT14 | 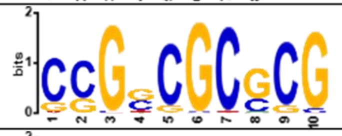 | 90.2 | 8.22e-5 |
| SP1/KLF12 | 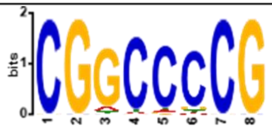 | 88.4 | 9.99e-5 |
| TAF1/HEN1 | 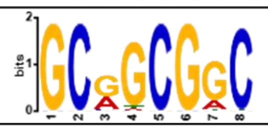 | 86.9 | 1.1e-4 |
| E2F6/KLF12 | 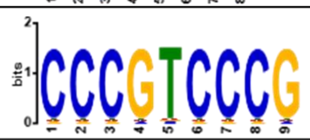 | 80.2 | 2.34e-4 |

**F**

#### Lost Super Enhancers

| TF | Consensus | TP% | p value |
|---|---|---|---|
| IRF1/2 | 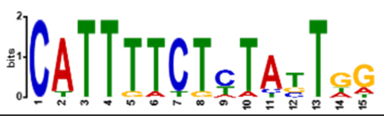 | 56.25 | 2.14e-12 |
| MEF2A/2B | 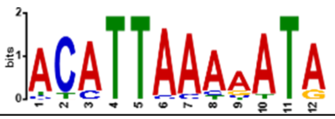 | 87.5 | 1.04e-11 |
| TBP | 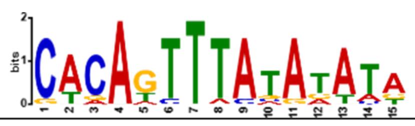 | 75 | 2.68e-10 |
| Tp53/63/73 | 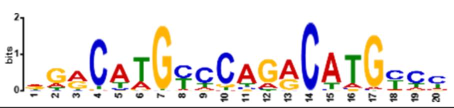 | 87.5 | 0.00165 |

**Fig S5**

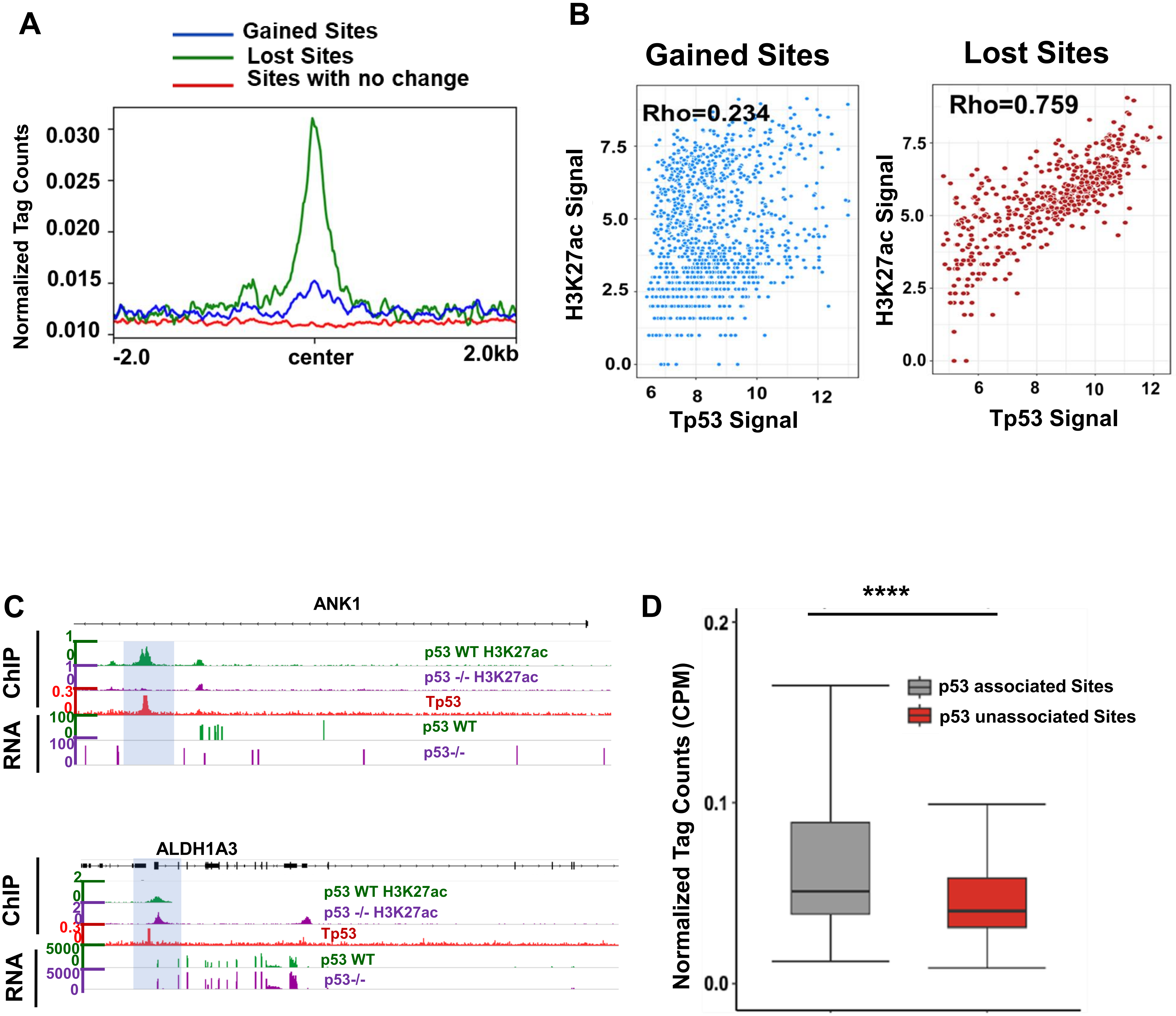
A
Gained Sites
Lost Sites
Sites with no change
Normalized Tag Counts
0.030
0.025
0.020
0.015
0.010
-2.0
center
2.0kb
B
Gained Sites
Rho=0.234
H3K27ac Signal
Tp53 Signal
Lost Sites
Rho=0.759
7.5
5.0
2.5
0.0
6
8
10
12
C
ANK1
ChIP
RNA
p53 WT H3K27ac
p53 -/- H3K27ac
Tp53
p53 WT
p53-/-
ALDH1A3
D
****
Normalized Tag Counts (CPM)
p53 associated Sites
p53 unassociated Sites
0.2
0.1
0.0

Fig S6

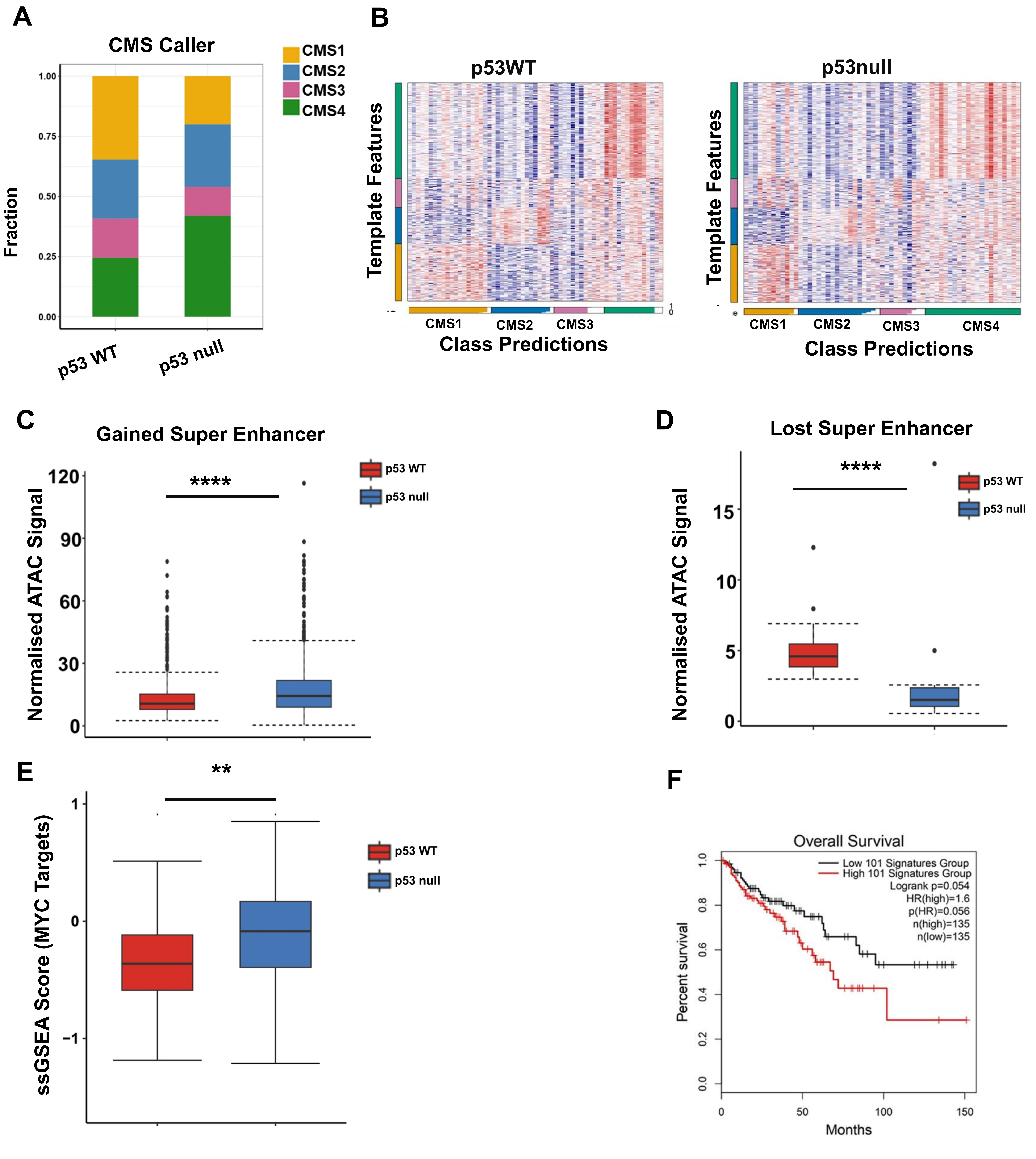
A
CMS Caller
CMS1
CMS2
CMS3
CMS4
Fraction
1.00
0.75
0.50
0.25
0.00
p53 WT
p53 null
B
p53WT
p53null
Template Features
CMS1
CMS2
CMS3
CMS4
Class Predictions
C
Gained Super Enhancer
****
p53 WT
p53 null
Normalised ATAC Signal
120
90
60
30
0
D
Lost Super Enhancer
****
15
10
5
0
E
**
ssGSEA Score (MYC Targets)
1
0
-1
F
Overall Survival
Low 101 Signatures Group
High 101 Signatures Group
Logrank p=0.054
HR(high)=1.6
p(HR)=0.056
n(high)=135
n(low)=135
Percent survival
1.0
0.8
0.6
0.4
0.2
0.0
0
50
100
150
Months

**Fig S1 ChIP-Sequencing of replicates confirm the global increase of H3K27ac due to loss of p53 in CRC cells**

**A.** Signal Intensity profile of H3K27ac in Replicate2 centered at +/-2kb of gained, lost and no change sites upon p53 loss in CRC cells. The profile represents an overall gain in H3K27ac upon loss of p53 consensus among the replicates.

**B.** Box plot representing distribution of normalized tag counts of H3K27ac at total gained and lost sites (+/-2Kb of center) upon p53 loss in CRC cells [p53 WT (red) and p53-/- (blue)] (Wilcoxon-Rank test was performed for the statistical analysis, p value> 0.05, ns; p value<=0.05, *; p value<=0.01,**; p value<=0.001,***). Gained H3K27ac sites showed significant higher levels of H3K27ac signals, while the lost H3K27ac sites showed significant lower levels of H3K27ac signals

**C.** Venn Diagram representing the overlap of H3K27ac ChIP-seq peaks at promoters between p53WT and p53-/- CRC cells. Increase in the number of active promoters sites upon loss of p53 in CRC cells

**D.** Venn Diagram representing the overlap of H3K27ac ChIP-seq peaks at enhancers between p53WT and p53-/- CRC cells. Increase in the number of distal enhancers upon loss of p53 in CRC cells.

**E.** Number of at promoters(red) and enhancers(blue) around the gained sites and lost sites respectively represented as stacked bar plot. Promoter-centric gain of H3K27ac sites upon p53 loss in CRC cells

**Fig S2 Alteration in H3K27ac levels is linked with gene expression dysregulation**

**A.** PCA plot representing distinct variation in the gene expression profile of p53 WT (red) and p53-/- CRC cells (blue).

**B**. Box plot representing distribution of gene expression profiles involved in p53 signalling in p53 WT (red) and p53-/- (blue) CRC cells (t-test was performed for the statistical analysis, p value> 0.05, ns; p value<=0.05, *; p value<=0.01, **; p value<=0.001, ***). Genes involved in the p53 signalling pathway are significantly downregulated with loss of p53.

**C.** Box plot representing distribution of gene expression profiles of MYC targets in p53 WT (red) and p53-/- (blue) CRC cells (t-test was performed for the statistical analysis, p value> 0.05, ns; p value<=0.05, *; p value<=0.01, **; p value<=0.001, ***). Genes involved in MYC targets are significantly upregulated with the loss of p53.

**D**. Heat map representing the number of differentially expressed upregulated and downregulated genes associated with gained and lost H3K27ac sites at the promoters. Number of genes correlates with gained and lost H3K27ac sites at the promoters.

**E.** Heat map representing the number of differentially expressed upregulated and downregulated genes associated gained and lost enhancer regions. The number of upregulated and downregulated genes correlates with the gained and lost H3K27ac sites at the enhancers respectively.

**F.** Positive enrichment score of MYC targets and E2F-targets associated with gained promoters and enhancers upon p53 loss as revealed through Gene set enrichment analysis (GSEA) on differential enrichment of gene signatures with p value <=0.05. Elevation of H3K27ac at regulatory elements upon loss of p53 correlates with upregulation of various MYC and E2F targets.

**G.** Negative enrichment of Interferon gamma response and complement pathways associated with lost promoters and enhancer sites upon p53 loss in CRC cells with p value <=0.05. Lost H3K27ac sites at regulatory elements correlates with down regulation of Interferon pathways.

**Fig S3 Distinct transcriptional regulators correlate with altered gene expression profiles and alterations in H3K27ac levels**

**A and B Table representing the differential motif analysis**

**A.** Percentage of H3K27ac peaks associated with different transcription factor and their respective p value around the gained sites as against the lost sites. CG rich motifs enriched at the gained H3K27ac sites.

**B.** Percentage of H3K27ac peaks associated with different transcription factor and their respective p value around the lost sites as against the gained sites. AT rich motifs enriched at the lost H3K27ac sites

**C.** IGV screenshot representing H3K27ac profiles in p53 WT (green) and p53-/- (purple) CRC cells integrated with the RNA tracks at the MAZ promoters (p53 WT CRC cells (green) and p53-/- CRC cells (purple)). Increased expression of MAZ is associated with gain of H3K27ac around the promoters.

**D.** Box plot representing distribution of gene expression profiles of High and low E2F target genes in p53 WT (red) and p53-/- (blue) CRC cells (student t-test was performed for the statistical analysis, p value> 0.05, ns; p value<=0.05, *; p value<=0.01, **; p value<=0.001, ***). High E2F targets corresponds with significant increase in expression profile.

**E.** IGV snapshots representing H3K27ac profiles in p53 WT (green) and p53-/- (purple) CRC cells integrated with the RNA tracks at SuV39H1 and EZH2 (p53 WT CRC cells (green) and p53-/- CRC cells (purple)). Increased expression of SuV39H1 and EZH2 are associated with gain of H3K27ac around the promoters and putative enhancers

**Fig S4 Oncogenic Super Enhancers are indirectly regulated by p53**

**A**. Box plot representing distribution of H3K27ac signal intensity profile around lost typical enhancers and lost super enhancers (Wilcoxon-rank test was performed for the statistical analysis, p value > 0.05, ns; p value<=0.05,*; p value<=0.01,**; p value<=0.001,***) in p53 WT and p53-/- CRC cells. Super-enhancers are associated with significantly higher H3K27ac levels as compared to typical enhancers.

**B.** Box plot representing gene expression distribution profile (log2 FPKM) around lost typical enhancers and super enhancers (t-test was performed for the statistical analysis, p value> 0.05, ns; p value<=0.05, *; p value<=0.01, **; p value<=0.001, ***) in p53 WT and p53 -/- CRC cells. Expression of genes associated with lost super-enhancers are higher as compared to typical enhancers

**C**. Gene set enrichment analysis representing positive enrichment score of MYC and E2F targets associated with gained super enhancers. Reprogramming of super-enhancers upon loss of p53 drives elevated expression of E2F and MYC targets.

**D**. IGV screenshot representing H3K27ac profiles in p53 WT (green) and p53-/- (purple) CRC cells integrated with the RNA tracks around the super enhancers of oncogenes TRA2B. Increased expression of E2F and MYC targets TRA2B are associated with gain of H3K27ac around the super-enhancers.

**E and F** Table representing the differential motif analysis **E.** Percentage of H3K27ac peaks associated with different transcription factor and their respective p value around the gained super-enhancer sites. Zinc finger nucleases (ZNFs) like transcription factors are enriched at gained super-enhancers.

**F.** Percentage of H3K27ac peaks associated with different transcription factors and their respective p value around the lost super-enhancer sites. IRFs are enriched at lost super-enhancers.

**Fig S5 Wild type p53 strongly correlates with the lost H3K27ac sites**

**A.** Plot Profile representing Tp53 signal (Naive) around gained(blue), lost(green) and no change sites(red). Tp53 strongly correlates with the lost H3K27ac sites

**B.** ScatterPlot representing correlation between Tp53 signal and H3K27ac signal at Gained(blue) and Lost (Red) sites. Tp53 shows higher significant positive correlation with H3K27ac around the lost sites.

**C.** IGV snapshots showing the changes of histone marks, p53 binding sites and gene expression in ANK1 and ALDH1A3. Decreased expression of ANK1 correlates with loss of H3K27ac. ANK1 was downregulated upon p53 loss. Increased expression of ALDH1A3 correlates with gain of H3K27ac. ALDH1A3 was upregulated upon p53 loss.

**D.** Box plot representing the distribution of p53 signal around the bound (grey) and unbound (red) H3K27ac sites, with higher p53 signal around bound sites as compared to the unbound sites (Wicoxon-rank test was performed for the statistical analysis, pvalue > 0.05, ns; p value<=0.05, *; p value<=0.01, **; p value<=0.001, ***). Tp53 signals are significantly higher around the sites associated with H3K27ac as compared to sites that are not associated.

**Fig S6 p53 null colon tumours are enriched with higher CMS proportion and increased accessibility at the gained oncogenic super-enhancer region.**

**A.** Stacked bar plot representing CMS classification in WT p53 and p53null COAD patients based on CMS classifier. Patients with p53 null mutation shows higher fraction of tumour population enriched with CMS4(green).

**B.** The consensus molecular subtypes (CMS) classification of CRC samples using R package CMScaller in p53 WT (left panel) and p53 null (right panel). p53null tumours has higher fraction of CMS4 subtypes as compared to p53WT.

**C.** Box plot representing the ATAC signals at the gained super-enhancers in p53WT (red) and p53 null tumours (blue) (t-test was performed for the statistical analysis, pvalue> 0.05, ns; p value<=0.05, *; p value<=0.01, **; p value<=0.001, **). p53 null tumours associates with higher accessibility at the gained super-enhancers as compared to p53WT tumours.

**D.** Box plot representing the ATAC signals at the lost super-enhancers in p53WT (red) and p53 null tumours (blue) (t-test was performed for the statistical analysis, pvalue> 0.05, ns; p value<=0.05, *; p value<=0.01, **; p value<=0.001, **). p53 null tumours associates with lower accessibility at the lost super-enhancers as compared to p53WT tumours.

**E.** Box plot representing distribution of ssGSEA score for MYC targets in p53 WT tumours(red) and p53 null tumours(blue). p53 null tumours associates with higher ssGSEA score for MYC targets as compared to p53WT tumour (t-test was performed for the statistical analysis, pvalue> 0.05, ns; p value<=0.05, *; p value<=0.01, **; p value<=0.001, **)

**F.** High median expression value of super enhancer genes associates with poor survival in TCGA-COAD (n=270) patients [HR =1.5 and p value <=0.1].
